# GeneMark-EP and -EP+: eukaryotic gene prediction with self-training in the space of genes and proteins

**DOI:** 10.1101/2019.12.31.891218

**Authors:** Tomáš Brůna, Alexandre Lomsadze, Mark Borodovsky

**Author notes:** Joint first authors (Tomáš Brůna, Alexandre Lomsadze).

## Abstract

We have made several steps towards creating a fast and accurate algorithm for gene prediction in eukaryotic genomes. First, we introduced an automated method for efficient *ab initio* gene finding, GeneMark-ES, with parameters trained in iterative *unsupervised* mode. Next, in GeneMark-ET we proposed a method of integration of unsupervised training with information on intron positions revealed by mapping short RNA reads.

Now we describe GeneMark-EP, a tool that utilizes another source of external information, a protein database, readily available prior to a start of a sequencing project. A new specialized pipeline, ProtHint, initiates massive protein mapping to genome and extracts hints to splice sites and translation start and stop sites of potential genes. GeneMark-EP uses the hints to improve estimation of model parameters as well as to adjust co-ordinates of predicted genes if they disagree with the most reliable hints (the -EP+ mode).

Tests of GeneMark-EP and -EP+ demonstrated improvements in gene prediction accuracy in comparison with GeneMark-ES, while the GeneMark-EP+ showed higher accuracy than GeneMark-ET. We have observed that the most pronounced improvements in gene prediction accuracy happened in large eukaryotic genomes.

## Introduction

One of major challenges of gene prediction in eukaryotes is finding an optimal way to combine sources of information extrinsic and intrinsic to genome of interest. External information could be transferred from RNA transcripts as well as from cross-species proteins derived from annotated genomes. Integration of transcript information, e.g. RNA-Seq reads, with *ab initio* gene prediction was implemented in several algorithms and software tools, e.g. AUGUSTUS (1), GeneMark-ET (2), EuGene (3,4), mGene.ngs (5). Also, a few other tools made use of protein sequences. Complexity of a task of leveraging cross-species protein sequence information for gene identification in a newly sequenced genome is growing with increase of evolutionary distance. Therefore, mapping a protein to genomic locus where a homologous protein is expected to be encoded was a subject for developing specialized tools known as tools for protein spliced alignment (e.g. currently available GeneWise (6), GenomeThreader (7), ProSplign (8), Spaln (9)). Beyond a single reference protein, a reference family of homologous proteins could be used to map elements of gene structure conserved in evolution, for instance, AUGUSTUS-PPX (10) uses protein profiles derived from conserved protein domains. Information about intron position, conserved in protein primary structures of multiple homologs was used in another tool, GeMoMa (11). Notably, an attempt to combine protein profiles with intron position profiles for refinement of predicted genes was made by yet another method GSA-MPSA (12).

Weakness of methods solely relying on mapping homologous proteins lies in the patchiness of the evidence they generate; a sizable fraction of a whole complement of genes may code for proteins with few or no orthologues. Another weakness is that protein spliced alignments become less accurate as the distance between the two species increases. Therefore, *ab initio* gene finders (e.g. GENSCAN (13), GeneMark.hmm (14), AUGUSTUS (15) or GeneID (16)) have been a necessary part of genome annotation tools and pipelines (e.g. GNOMON (17), PASA (18) and Ensembl (19)).

Application of *ab initio* algorithms for genome wide eukaryotic gene prediction was for long time hampered by the need of tedious and time-consuming training. To address this issue we have earlier developed an *ab initio* gene finder GeneMark-ES (20,21) with model parameters estimated by iterative unsupervised training. This algorithm did not require expert based training or hints for building a training set. GeneMark-ET (2) was developed to make GeneMark-ES able to integrate into training process available transcript information, raw RNA-Seq reads spliced aligned to genome in question.

Here we describe GeneMark-EP, an algorithm and software tool that integrated into training information extracted from a reference set of cross-species protein sequences. To generate protein hints for a given genomic locus we first identify a set of proteins, homologous to the protein likely encoded in the genomic locus. Then a specialized pipeline, ProtHint, computes the hints, a set of mapped splice sites (intron borders) and translation start and stop sites along with the scores characterizing hint confidence. The most reliably determined elements of spliced alignment could be used to directly identify elements of exon-intron structures, this mode of algorithm execution with direct gene structure correction we call GeneMark-EP+.

A key question is how to find optimal method of hint incorporation into the *ab initio* algorithm. Unsupervised training implemented in GeneMark-ES carries a risk of convergence to a biased set of model parameters. On the other hand, giving too much weights to protein hints may generate parameters dictated by a narrow set of conserved genes and proteins (22). By design, the GeneMark-EP algorithm combines strong features of both methods: i/ ability of unsupervised iterative training of an *ab initio* gene finder to create a set of training sequences with a size beyond reach of conventional supervised training and ii/ ability to correct model parameters as well as (the EP+ mode) structures of newly discovered genes with the hints derived from homologous cross-species proteins. The new method falls into category of gene prediction methods with semi-supervised training.

## Materials

For assessment of GeneMark-EP as well as ProtHint accuracy we selected annotated genomes from diverse clades: fungi, worms, plants, insects, and vertebrae (Table 1). The genomes length varied from under 100 Mb *(Neurospora crassa)* to more than 1.3 Gb *(Danio rerio).* With exception of *Solanum lycopersicum,* a species representating large genome plants important for economy, all selected species are model organisms whose genomes presumably have high quality annotation. To assess accuracy of gene prediction made for model species, we compared genes predicted and annotated on a whole genome scale. In case of *S. lycopersicum* we used a limited set of genes, validated by available RNA-Seq data. In all genomic datasets, contigs not assigned to any chromosome were excluded from the analysis as well as genomes of organelles.

**Table 1:**
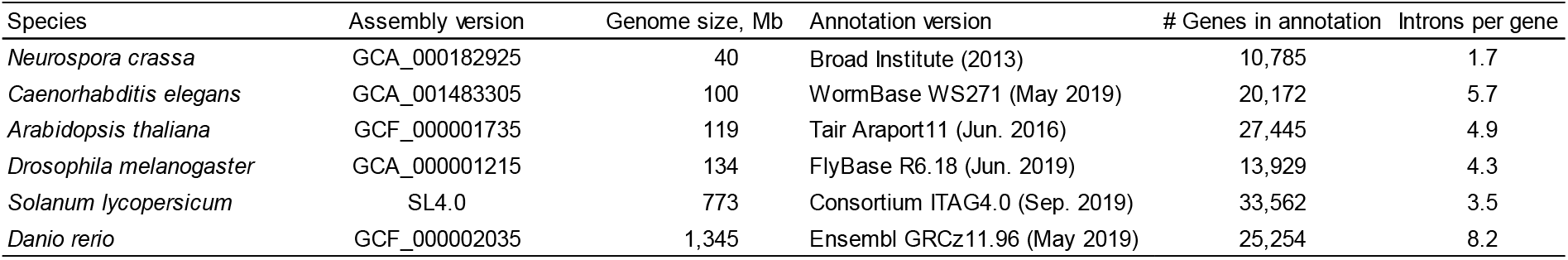
Genomes used for assessment of GeneMark-EP and GeneMark-EP+ performance. Introns per gene values were computed with respect to the whole gene number, including singleexon genes.

| Species | Assembly version | Genome size, Mb | Annotation version | # Genes in annotation | Introns per gene |
| --- | --- | --- | --- | --- | --- |
| *Neurospora crassa* | GCA_000182925 | 40 | Broad Institute (2013) | 10,785 | 1.7 |
| *Caenorhabditis elegans* | GCA_001483305 | 100 | WormBase WS271 (May 2019) | 20,172 | 5.7 |
| *Arabidopsis thaliana* | GCF_000001735 | 119 | Tair Araport11 (Jun. 2016) | 27,445 | 4.9 |
| *Drosophila melanogaster* | GCA_000001215 | 134 | FlyBase R6.18 (Jun. 2019) | 13,929 | 4.3 |
| *Solanum lycopersicum* | SL4.0 | 773 | Consortium ITAG4.0 (Sep. 2019) | 33,562 | 3.5 |
| *Danio rerio* | GCF_000002035 | 1,345 | Ensembl GRCz11.96 (May 2019) | 25,254 | 8.2 |

We used OrthoDB v10 protein database (23) as an all-inclusive source of protein sequences. Still, for generating protein hints for particular species we used subsets of OrthoDB: plant proteins for gene prediction in *Arabidopsis thaliana,* arthropod proteins - for *Drosophila melanogaster,* etc. (Table 2).

**Table 2:** Characteristics of the OrthoDB v10 taxonomical space for each of the species we tested. The number of species is naturally the largest in the Kingdom section of the database. *For tests in the genus-, family-, and order-excluded modes for D. *melanogaster* and D. rerio, the phylum was used as the largest set of reference proteins.

| Number of species in the same taxonomical unit | Genus | Family | Order | Class | Phylum | Kingdom | OrthoDB root used for tests | # of proteins in the root |
| --- | --- | --- | --- | --- | --- | --- | --- | --- |
| *Neurospora crassa* | 0 | 1 | 7 | 96 | 364 | 548 | Fungi | 5,850,648 |
| *Caenorhabditis elegans* | 2 | 2 | 4 | 5 | 6 | 447 | Metazoa | 8,266,016 |
| *Arabidopsis thaliana* | 1 | 7 | 9 | - | 99 | 116 | Plantae | 3,510,742 |
| *Drosophila melanogaster* * | 19 | 19 | 55 | 147 | 169 | 447 | Metazoa | 8,266,016 |
| *Solanum lycopersicum* | 1 | 9 | 10 | - | 99 | 116 | Plantae | 3,510,742 |
| *Danio rerio* * | 0 | 4 | 4 | 49 | 245 | 447 | Metazoa | 8,266,016 |

As an additional test set we used annotation of major protein isoforms available in the APPRIS database (24); this assessment was done for *C. elegans*, *D. melanogaster*, *and D. rerio* (Table S1). Arguably, accuracy of prediction of major isoforms is of significant interest, since in a gene locus the major isoform was observed to be expressed in higher volume than other (minor isoforms) (24).

## Methods

### Integration of genomic sequence patterns and protein homology into gene prediction

The GeneMark-EP, -EP+ algorithm goes step-by-step through the following tasks: i/ selection of genomic regions, *seed regions,* containing gene candidates *(the seed genes);* ii/ identification for each seed gene a set of homologous proteins iii/ constructing splice alignments of homologous proteins to each seed region and generating hints for exon-intron structures; iv/ running itereative semi-supervised training with selection of most reliable (*anchored*) elements of predicted genes in each iteration; v/ final gene prediction without (-EP mode) or with additional option (-EP+ mode) to enforce high confidence hints in predicted exon-intron structures (Fig. 1).

**Figure 1:**
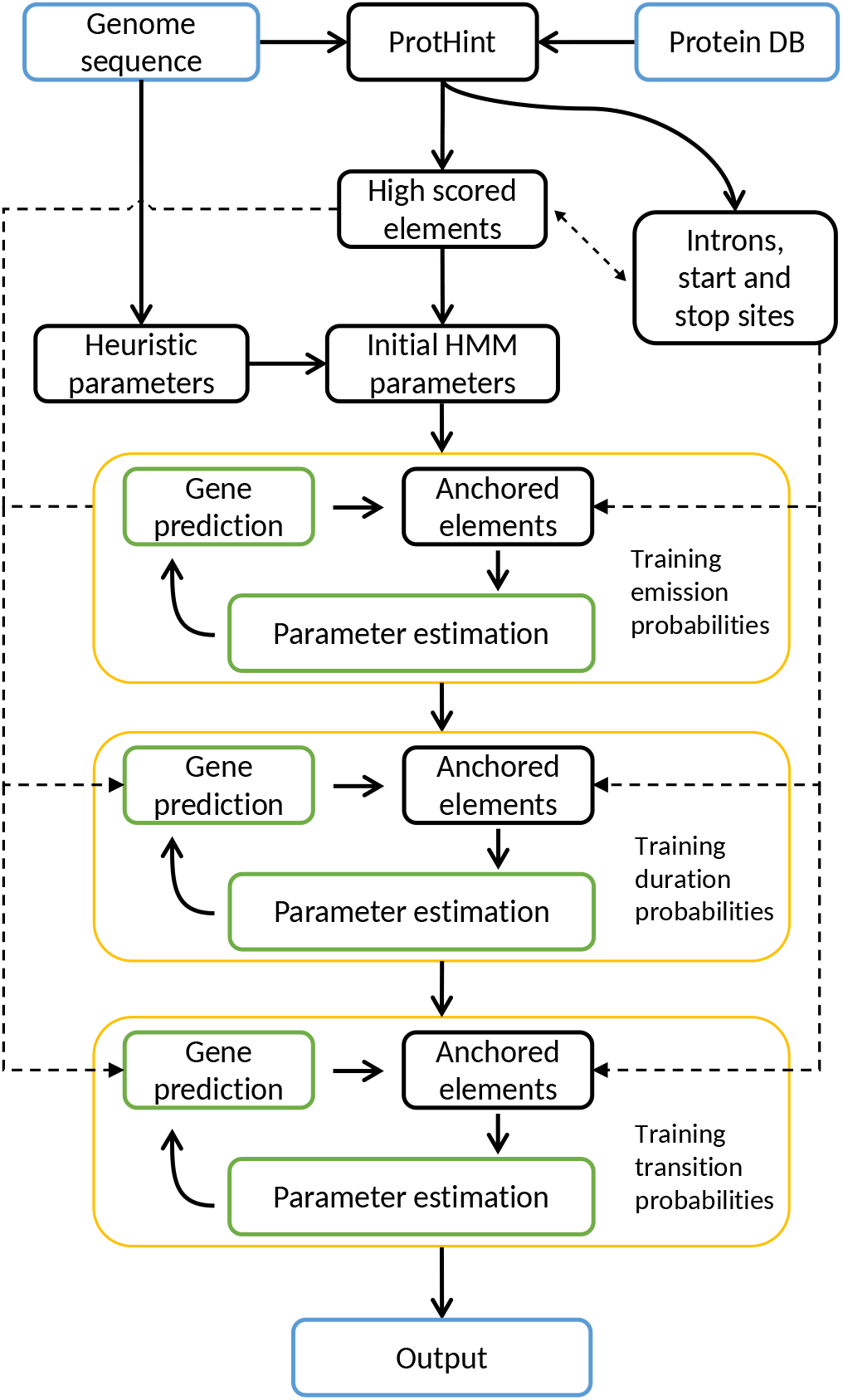
A flowchart of the GeneMark-EP, EP+ iterative training.

The tasks i/-iii/ are devoted to generating protein hints and are solved by the ProtHint pipeline (Fig. 2). To determine *seed regions* within a long genomic sequence (task i/) we run unsupervised training of parameters of statistical models by GeneMark-ES (20) and generate *ab initio* gene predictions. To create a *seed region,* each predicted gene, a *seed gene,* is expanded upstream and downstream by adding 2,000 nt margins. To identify proteins homologous to a *seed protein,* task ii/, we run DIAMOND similarity search (25) with a *seed protein* as a query against a protein sequence database (e.g. a section of OrthoDB). A set of proteins with statistically significant hits define a set of target proteins presumed to be homologous to the query, the seed protein. Task iii/ is to generate spliced alignments of multiple protein targets to the seed region (done by either Spaln (9) or ProSplign (8)) and to process the results of alignments in order to infer elements of exon-intron structures (introns, splice sites, translation starts and stops) characterized by reliability scores. Mapped gene elements with reliability scores exceeding certain thresholds are designated as high-confidence hints. Final tasks (iv) and (v) correspond to training and prediction steps of GeneMark-EP and -EP+. At these steps we use the hints to exon-intron structure co-ordinates as an input to an expectation-maximization type algorithm that finds models of compositional patterns of protein-coding and noncoding regions simultaneously with the most likely parse of genomic sequence into coding and non-coding regions.

**Figure 2:**
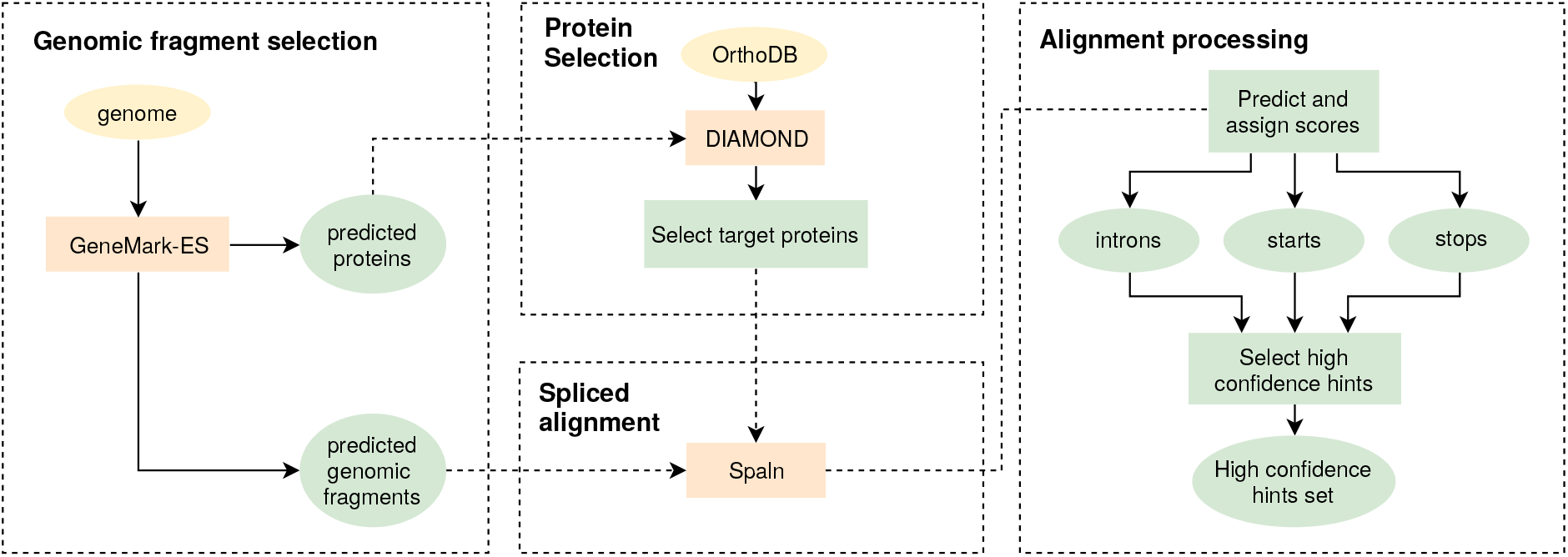
An overview of the ProtHint pipeline.

Iterative training of the GeneMark-EP statistical models (tasks iv and v) works as follows. In the first iteration full length introns mapped by ProtHint with scores exceeding a stringent threshold (high confidence elements) are used to estimate parameters of splice sites models as well as branch point site models (particularly important for intron models of fungal genomes). The splice site models together with the heuristic models of protein-coding and non-coding regions make a complete set of models of a semi-Markov HMM (20). The models are used in the first run of the Viterbi algorithm (see (14)) that generates a maximum likely parse of genomic sequence into coding and non-coding regions, the parse delineating the first set of genes predicted by GeneMark-EP. Next, we analyze available data to make updated training sets and re-estimate model parameters. We compare co-ordinates of exons predicted by GeneMark.hmm and intron hints determined by ProtHint within the *seed* regions. This comparison leads to selection of *anchored* elements, the exons with at least one splice site identified by both GeneMark.hmm and ProtHint. A set of anchored exons along with a set of predicted single exon genes (with length > 800nt) comprise an updated training set for the three-periodic Markov chain model of protein-coding region (26). Sequences of introns bounded by two anchored splice sites as well as intergenic sequences bordered by anchored terminal and initial exons of adjacent genes (Fig. 3) are used for updating parameters of the non-coding region model. The set of updated models is used by the Viterbi algorithm to generate a new set of predicted genes. A new update of anchored elements and the next round of parameter re-estimation follows.

**Figure 3:**
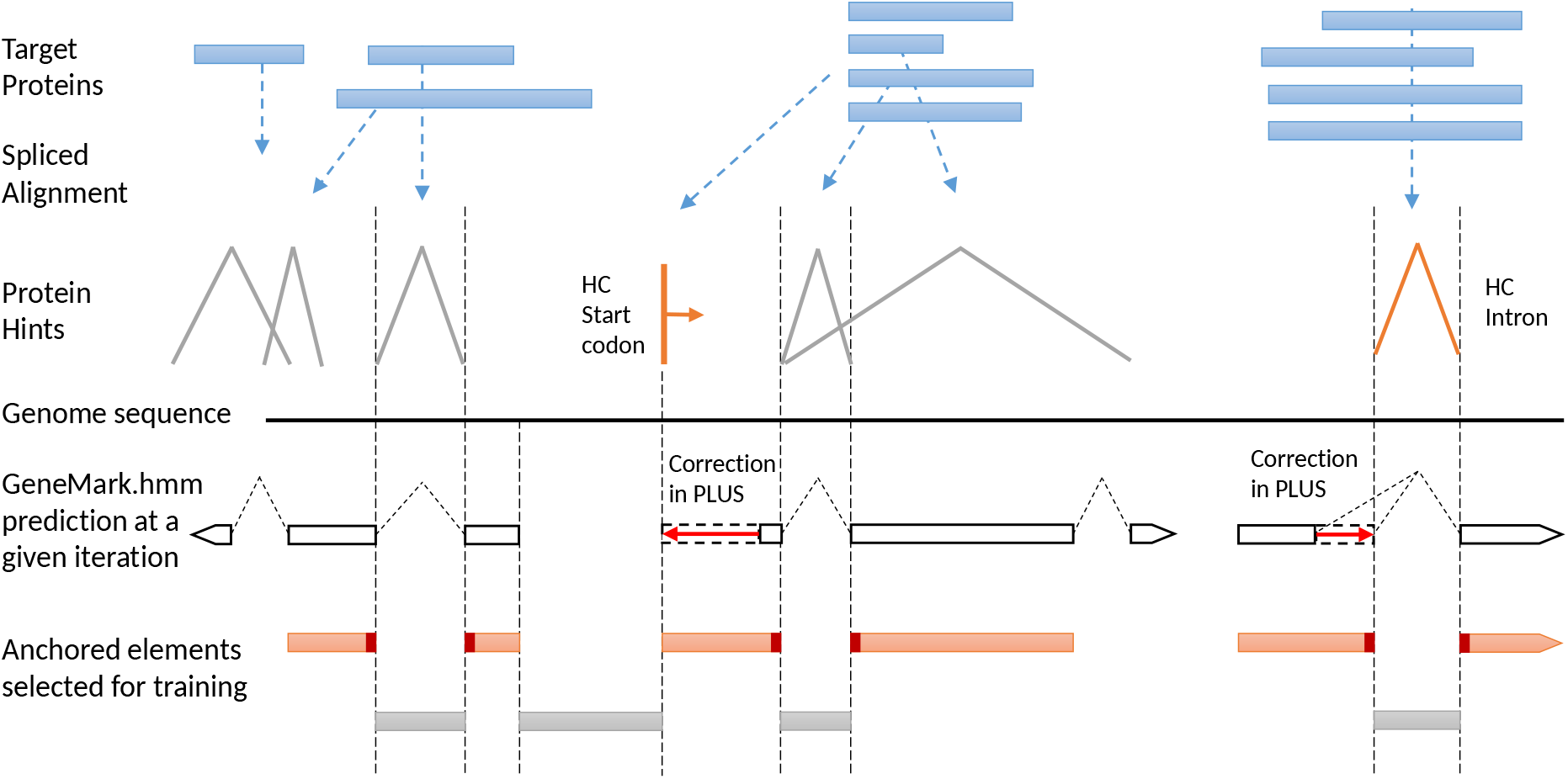
Selection of anchored elements for GeneMark-EP+ training with enforcement of High-Confidence (HC) hints.

Several probability distributions used in GeneMark-EP, such as length distributions of exon, intron and intergenic regions, are initially defined as uniform ones. More accurate estimation of these distributions is done in subsequent steps of iterative training (Fig. 1). Also, in the later steps we estimate parameters of the three-phase models of splice sites indexed by a nucleotide position after which the intron divides a codon triplet. Prior to the final iteration we update estimates of the HMM transition probabilities that affect frequencies of genes with given number of introns. In experimental runs for genomes with different length we have verified that seven iterations were sufficient for GeneMark-ES and six iterations for GeneMark-EP and -EP+ to reach convergence in co-ordinates of predicted genes and values of model parameters.

Gene predictions made in the final iteration are reported as output of GeneMark-EP. Running the Viterbi algorithm could be done with enforcing high confidence elements mapped by ProtHint. Particularly, it is done by modifying components of the object function of the Viterbi algorithm associated with chosen hidden states. The sites that must be enforced receive high values of objective function to ensure their addition to a path selected by the optimization algorithm seeking the maximum value of the log Viterbi objective function. This mode of gene prediction produces the GeneMark-EP+ output.

Note that GeneMark-ES, -ET, -EP, -EP+ algorithms are designed to predict non-overlapping genes with no alternative isoforms. This design suits the paradigm that each gene locus encodes a major (expressed in most tissues) protein isoform and majority of alternative transcripts have differences in situated in 5’ and 3’ UTRs (24).

### ProtHint: generating footprints (hints) of multiple homologous proteins for a genomic locus

#### General logic

The ProtHint role (Fig. 2) in GeneMark-EP, - EP+ is two-fold. This pipeline generates two sets of protein hints. The smaller one, the set of high confidence hints, includes hints with high scores that ensure their high specificity. The larger one includes hints that have scores exceeding a liberally set threshold, thus these hints have lower specificity but larger sensitivity. In the process of hint generation ProtHint takes a *seed protein* and uses it as a query in similarity search for homologs of a true protein presumably encoded in the seed region. Next, ProtHint constructs spliced alignments of the detected homologs (target proteins) to the seed region. The whole set of multiple spliced alignments is then processed together to identify protein hints, mapped co-ordinates of the candidate splice sites as well as translation start and stop sites. Hints scoring system is discussed in detail in Supplementary Materials.

Technically, for a given *seed pro*tein, ProtHint runs DIAMOND (25) against a relevant section of the OrthoDB database and retains in the output up to 25 target proteins (with hit E-value better than 0.001). Next, the target proteins are spliced aligned by Spaln (9) back to the seed region. Notably, the hints are defined by ProtHint processing of the Spaln raw pairwise alignments instead of using annotation of exons in the Spaln output. Rather frequently, multiple target proteins aligned to the original seed region may map out one and the same sequence fragment as an intron. Such an outcome would define an intron hint with a higher confidence than if an intron candidate is mapped only once.

#### Score system for introns

As described above an expected evolutionary conservation between primary structures of target proteins and a protein encoded in the seed region has to be quantified and used for accurate identification of a new gene. To facilitate this quantification, we define three types of scores for exons and introns (AEE, IBA and IMC, see below) and two types of scores for candidate gene starts and stops (SMC and BAQ, see below).

##### Alignment of Entire Exon

(AEE) score is defined as a score of the Spaln (or ProSplign) alignment of exon translation and a target protein (for more details see Supplementary Materials).

##### Intron Borders Alignment

(IBA) score is computed from kernel modified alignment scores of two adjacent exons with larger weights given to parts close to the donor and acceptor splice sites. An IBA score (within a window of length *w*, being 10 amino acids by default) is computed as follows.

For downstream (and upstream) exon defined in the Spaln spliced alignment we compute *S_d_* (and *S_u_*) as

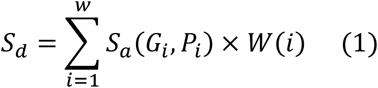

Here *S_a_*(*G_i_, P_i_*) is a substitution score defined for target protein amino acid *P_i_* and a codon defined amino acid *G_i_*; *W*(*i*) is the weight function. For instance, for a downstream exon *S_d_*:

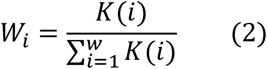

where *K*(*i*) is the kernel value for position *i* counting in codons from an acceptor site. In a linear kernel:

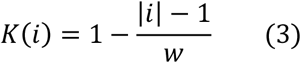

Then we take a geometric mean of values of *S_d_* and *S_u_*.

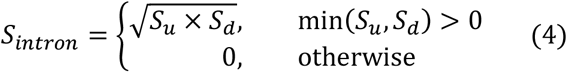

Finally, the IBA score is obtained by normalizing the *S_intron_* score into 〈0,1〉 range: IBA score = *S_intron_* / max(*S_a_*), where max(*S_a_*) is a maximum score among elements of the BLOSUM62 matrix.

##### Intron Mapping Coverage

(IMC) score is a count of how many times a given intron was exactly mapped by spliced alignments of target proteins. The IMC score is computed only from the set of the introns which IBA score exceeds certain level.

Notably, instead of all introns with identical coordinates and different IBA scores related to different target proteins we deal with a single intron characterized by the maximum of individual IBA scores among all collapsed introns.

#### Application of the intron scores

For a particular seed region, we use the following method to define hints to exonintron structure from a set of spliced alignments of target proteins:

First, we select introns whose two adjacent exons have scores AEE ≥ E_t_, where E_t_ is a chosen threshold. For E_t_ =25, in a modeling on known genomes, we observed relatively high Sn value of the candidate introns (Fig. S1). Further increase of *E_t_* eliminated true introns while not significantly improving Sp value.

Next, to reduce number of false positives in the obtained set of introns we selected a subset with the IBA score > I_t_ where I_t_ is another chosen threshold. Our modeling has shown an increase in the Sp value of the candidate introns for I_t_ = 0.1 that occurred without noticeable change in Sn (Fig. S1).

Thus identified sets of introns for all the seed regions represents a set of ***all*** mapped introns; it is used as external evidence to generate *anchored introns* for GeneMark-EP training steps as described above.

Finally, within the set of ***all*** mapped introns we select a narrower set of *high-confidence introns.* These introns must have canonical GT-AG splice sites, an IMC score ≥ 4, and an IBA score ≥ 0.25 (Figs. 4, S2).

**Figure 4:**
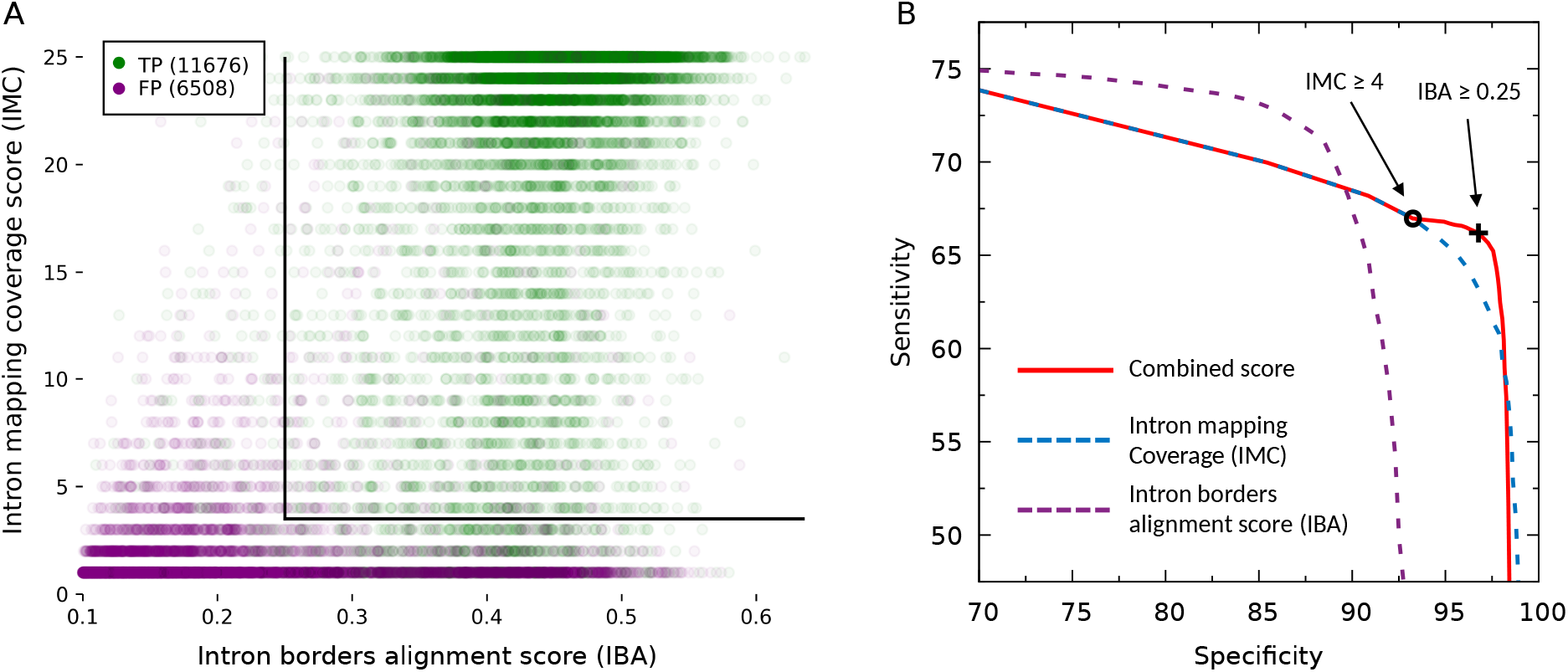
ProtHint intron processing in case of *N. crassa.* Introns were generated by spliced alignments of target proteins from species beyond *Neurospora* genus. (a) Distribution of the score vectors (IBA, IMC) of true positive (green) and false positive (purple) introns. The black lines represent cutoffs at IMC = 4 and IBA = 0.25. Total numbers of false and true positives are shown in the upper left corner. (b) Sn and Sp of intron sets selected by thresholds on IBA score and IMC score. IMC score is computed for introns which have IBA score ≥ 0.1 and exon AEE score ≥ 25. The red curve represents the following. The left branch of the curve reflects (Sp, Sn) values of the sets of introns selected by using IMC threshold from 0 to 4. The one with the IMC threshold = 4 is recorded as set A – the set corresponding to the black circle in the red curve. Then, the right branch of the curve reflects (Sp, Sn) of the set of introns generated by applying to set A an IBA score threshold changing from 0 to 0.25 and up to 1.0. Set B corresponds to the black cross in the red curve, introns in this set have IMC ≥ 4 and IBA ≥ 0.25. Separate curves for IMC score change (dashed blue) and IBA score change (dashed purple) are shown as well.

We use high-confidence introns to estimate initial parameters of the GeneMark-EP intron model. Besides, these introns are enforced in prediction step of the GeneMark-EP+ mode.

#### Score system for translation starts and stops

Similarly, to scores introduced for intron hints generation, we define a ***Border Alignment Quality*** (BAQ) score for translation starts and stops. This score is computed for *w* amino acids downstream (upstream) of start (stop) codon, weighted by a kernel dependent function (Eq. 1).

The second type of score is the ***Site Mapping Coverage*** (SMC) score. This score is a count of N-terminals (C-terminals) of target proteins aligned to a particular start (stop) codon position of a candidate gene. The SMC scores are computed only from the sets of initial (terminal) exons whose BAQ scores exceed certain level.

If a set of target proteins for a given seed region generates footprints situated more upstream than others, alternative start candidates situated downstream are removed from consideration (Fig. S3, details in Supplementary Materials). We have observed that using these rules leads to increase in the hints accuracy (Table 3, S2).

**Table 3:**
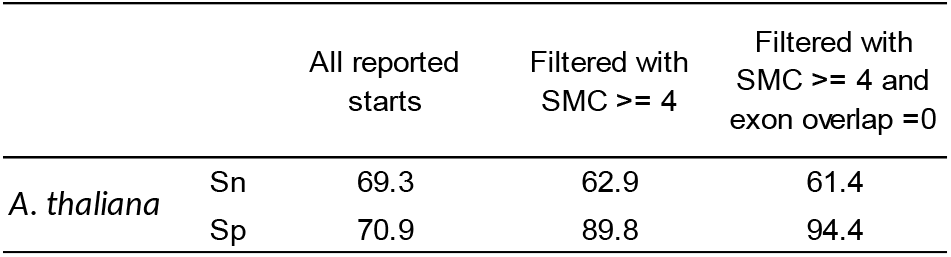
Case of *A. thaliana.* Sensitivity and specificity of all gene start hints created by ProtHint as well as of the high-confidence start hints. High specificity was achieved with filtering by SMC scores as well as by removal of candidate starts overlapped by at least one target protein which suggested an alternative start upstream. Sensitivity was defined with respect to a full complement of starts, including alternative ones as given in annotation. The numbers were generated in tests with reference proteins from species outside a relevant genus. Results for all test species are shown in Table S2.

#### Application of BAQ and SMC scores

All over, selection of a set of ***all*** translation start and stop hints is done by the following method:

A start codon candidate is an ATG codon present in a mapped initial exon and aligned to N-terminal methionine in a target protein; a stop codon candidate is a stop codon in a mapped terminal exon.

A candidate initial (terminal) exon containing candidate gene start (stop) should have AEE score ≥ 25 and BAQ score for candidate start (stop) codon ≥ 0.

To select a subset of *high-confidence hints* we choose stop codon candidates with SMC score ≥ 4 as well as start codon candidates with SMC score ≥ 4 *and* no overlap by longer target proteins (Fig. S3). The set of *high-confidence hints* to translation starts and stops is used to estimate parameters of GeneMark-EP models of translation initiation and termination sites. Also, the *high-confidence hints* are directly enforced in prediction step of GeneMark-EP+.

### Do introns mapped by ProtHint tend to occur in gene regions coding for conserved domains?

To address this question we use the following procedure. Annotated genes are translated to proteins and used as queries in RPS-BLAST (27) to search (E-value =0.01) against NCBI Conserved Domains Database (CDD) (28). Results of the RPS-BLAST searches are processed with *rpsbproc* utility (28) to generate a map of conserved domains for each RPS-BLAST query. Finally, coordinates of the conserved domains are mapped back to a *seed region* of genomic DNA and compared with the ProtHint output to find out how many introns are mapped into regions coding for conserved domains. We conducted this analysis for genes of *D. melanogaster*, *C. elegans*, and *D. rerio* genomes annotated in the APPRIS database (24) as genes coding for principal protein isoforms (see Results).

### Assessment of GeneMark-EP, EP+ gene merging and gene splitting errors

Gene merging and splitting errors are expected to be reduced by the use of homologous protein derived hints to gene translation starts and stops. This expected improvement in prediction accuracy of GeneMark-EP+ could be more accurately observed on properly prepared test sets. Prior to evaluation of gene splitting we had to exclude from the test sets: i/ genes fully overlapping shorter genes present inside introns in any strand; ii/ genes with larger isoforms combining or including shorter alternative components (Fig. S4); iii/ genes with introns longer than 10,000nt (the default maximum intron length). For genes with annotated multiple alternative isoforms we used the longest one as a representative. Prior to evaluation of gene merging overlapping genes present in annotation (e.g. a gene within an intron) were merged into a single gene in order to exclude such cases from being counted as merged genes.

## Results

We have compared gene prediction accuracy of GeneMark-EP, -EP+ with accuracy of GeneMark-ES and GeneMark-ET. In addition, we made an accuracy assessment of ProtHint. We worked with genomes of six species *N. crassa, C. elegans*, *A. thaliana*, *D. melanogaster*, *S. lycopersicum* and *D. rerio* (Table 1). Genomes of model organisms (all the species but *S. lycopersicum*) expected to have sufficiently accurate annotation. In such cases, we made comparisons between predicted and annotated gene co-ordinates on whole genome scales. In case of *S. lycopersicum* we built a test set of genes validated by RNA-Seq data.

In genomes of *C. elegans*, *A. thaliana*, *D. melanogaster*, and *D. rerio* were excluded from comparisons all regions of annotated pseudogenes. Also, in case of *D. rerio* we excluded annotated partial exons (ubiquitous in this genome) from exon level accuracy assessment, while we computed the gene level sensitivity only for genes having complete alternative annotated transcripts.

We used OrthoDB v10 (23) as a source of protein sequences partitioned into relevant taxonomic divisions; particularly, we used plant division for *A. thaliana*, arthropod division for *D. melanogaster*, etc. (Table 2).

A principal feature of the new method is use of *multiple homologous proteins* for hints generation. In a practical application to a novel genome, evolutionary distances *from encoded proteins of interest to sets of homologs* within already known protein universe could vary significantly depending on having genomes of close or moderately close species sequenced and annotated or not. To model these variations in our tests, we introduced restrictions on how evolutionary close the target proteins could be to a given query. These restrictions were implemented by removing from the protein database: i/ proteins encoded in the genome of a given species; ii/ - from all species from the same subgenus; iii/ - from the same genus; iv/ - from the same family; v/ - from the same order; vi/ - from the same phylum. Notably, distributions of numbers of species within a genus, family, etc. defined by a given species are species specific (Table 2).

### Assessment of accuracy of GeneMark-EP, -EP+

For each species (Table 1) we determined how the accuracy of GeneMark-EP, -EP+ at gene level (Fig. 5) and exon level (Fig. S5) depended on the choice of a set of reference proteins. The pattern of accuracy change at *gene level* (Fig. 5) was similar to the one observed at *exon level,* therefore, we show the results of accuracy assessment at gene level in the main text while the results for exon level are provided in Supplementary Material (Fig. S5). Even more details on accuracy assessment of GeneMark-EP (running without enforcement of high-confidence hints) and of GeneMark-EP+ are given in Supplementary Materials (Table S3).

**Figure 5:**
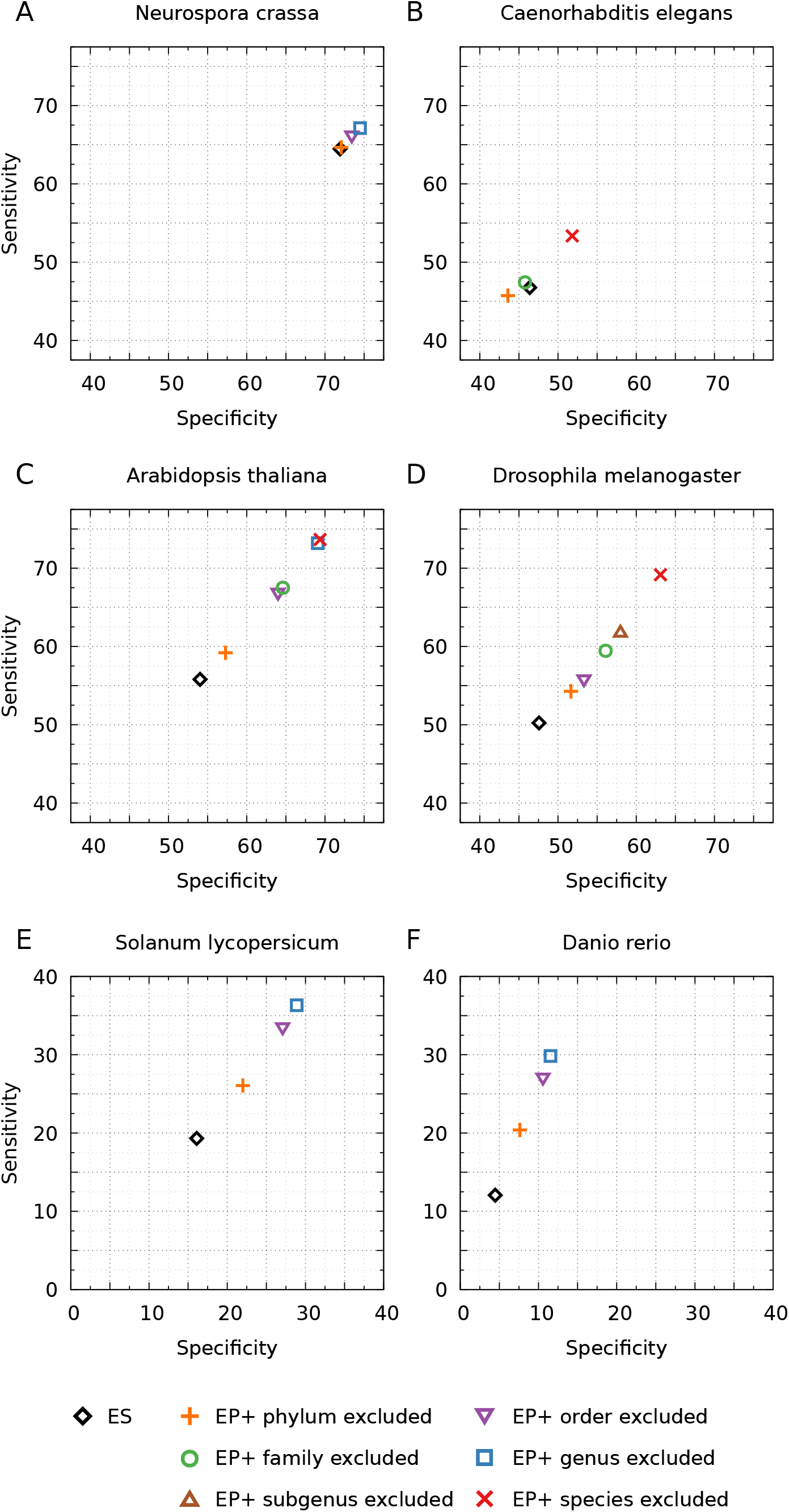
Comparison of GeneMark-ES and GeneMark-EP+ accuracy on gene level. Accuracy of GeneMark-EP+ is shown for cases when ProtHint works with different size sets of reference OrthoDB proteins: from the largest (only proteins from the same species are excluded) to the smallest (proteins of the whole phylum excluded). A gene prediction is considered to be correct if it matches one of the annotated isoforms. For *D. rerio* gene level Sn was computed only with respect to complete genes.

We present results for the three groups of genomes: fungal genomes, compact eukaryotic genomes and large eukaryotic genomes.

#### Fungal genomes

*N. crassa*. Accuracy of GeneMark-ES was high, as it has been typical for fungal genomes (21). Even with hints originated from the largest set of reference proteins, those outside genus/order, GeneMark-EP+ improved Sn value of GeneMark-ES only by ~2% (Fig. 5a). With smaller set of more remote reference proteins that originated from the species outside fungal phylum, the accuracy of GeneMark-EP+ matched the accuracy of GeneMark-ES (Fig. 5a). This result went in line with previous observations that GeneMark-ES was highly efficient *ab initio* gene finder for fungal genomes (21). We observed earlier that for fungal genomes a support of training with data from splice-aligned RNA-Seq reads makes accuracy of GeneMark-ET at best slightly higher than one of GeneMark-ES.

#### Compact eukaryotic genomes

*C. elegans*, *A. thaliana*, and *D. melanogaster*. When GeneMark-EP+ used the largest set of reference proteins (just without proteins from the same species) we saw for *A. thaliana* and *D. melanogaster* an improvement by ~20% in both Sn and Sp in comparison with GeneMark-ES (Figs. 5c,d). As target proteins were coming from larger and larger evolutionary distances, the accuracy did steadily decrease. When target proteins were selected outside the same phylum there was an increase by only 5% in gene level Sn and Sp in comparison with GeneMark-ES. For *C. elegans*, when the set of reference proteins excluded just proteins of the same species, GeneMark-EP+ improved the accuracy of -ES by ~6% (Fig. 5b). We observed almost no difference between GeneMark-EP+ and GeneMark-ES when the reference proteins were only from species outside the *C. elegans* family and a slight decrease in accuracy (by ~2%) for reference proteins outside of the taxonomical phylum. Notably, the gene level accuracy for *C. elegans* was lower than for other species with compact genomes.

#### Large eukaryotic genomes

*S’. lycopersicum* and *D. rerio*. The gene level accuracy of GeneMark-ES was low for these genomes (between 5% and 20%). GeneMark-EP+ improved the accuracy for *S. lycopersicum* by ~15%, when it used a protein reference set from species outside of the tomato genus or order (Fig. 5e). For *D. rerio,* having a reference set of proteins without ones from same genus or the same order as *D. rerio* led to Sn and Sp improvement by ~20% and ~5%, respectively (Fig. 5f). However, the improvements were twice as low when reference proteins were available only outside the *S. lycopersicum* or *D. rerio* phylum.

Relatively low gene prediction accuracy in large genomes could be partially attributed to incorrect and/or incomplete gene annotations. Therefore, we made additional effort to refine test sets in *D. rerio* and *S. lycopersicum* by selecting genes supported by RNA-Seq data.

We observed that if annotated genes of *S. lycopersicum* genome were supported by RNA-Seq, they were significantly better predicted by GeneMark-EP+ (Table S4). To generate intron hints from RNA-Seq we used VARUS (29). We divided annotated tomato genes into two groups: a/ genes with all introns predicted by VARUS and b/ all other genes. GeneMark-EP+ sensitivity (for a GeneMark-EP+ run having reference proteins outside the *S. lycopersicum* genus) was by 40% better in set (a) than in set (b), on gene, exon as well as intron levels. It is important to emphasize that RNA-Seq information was not used in GeneMark-EP+. Sensitivity defined for the set of introns mapped by ProtHint was also better in set (a) by ~40% (Table S4).

We already mentioned that *D. rerio* annotation had many partial exons that in turn would be parts of incomplete transcripts. We evaluated exon level Sn separately for exons within complete and incomplete transcripts (Table S5) and observed 75.1% exon Sn in ‘complete’ group vs 67.6% in ‘incomplete’ group. Similarly, gene level sensitivity was better by 6% in predicting genes with complete transcripts compared to all genes (Table S5).

All over, we observed that for majority of the considered species, the accuracy of GeneMark-EP+ was better than accuracy of GeneMark-ES, regardless of how large of a set of reference proteins was used for spliced alignments (Table S3, Figs. 5, S5). For the fungal genome, *N. crassa,* an improvement was negligible due to ability of GeneMark-ES deliver high accuracy for fungal genomes; we also observed a small decrease of accuracy in the *C. elegans* test with phylum excluded reference set of proteins (Fig. 5).

### Comparison with GeneMark-ET

In addition, we compared GeneMark-EP with GeneMark-ET [2] which uses RNA-seq short reads to provide external information (hints to intron co-ordinates) to select anchored gene elements for the GeneMark-ET algorithm parameter estimation. GeneMark-ET does not have an ‘-ET+’ mode in which predictions are directly guided by high confidence hints. We ran GeneMark-ET with hints to co-ordinates of introns mapped by VARUS (30) from RNA-Seq reads. VARUS automatically sampled, downloaded and aligned reads from NCBI’s Sequence Read Archive (SRA) with time stamp 01/22/2020 (31). The time stamp is important for the results reproduction, since the VARUS outcome depends on the amount of RNA-Seq data deposited to SRA. As one could see (Table S3) the accuracy of GeneMark-ET with training guided by hints derived from mapped RNA-Seq reads is very close to the accuracy of GeneMark-EP with training guided by hints derived from mapped proteins.

To get an idea to which extent a whole complement of genes is covered by hints originated from protein or transcript data, we determined fractions of genes with at least one hint or at least with one high-confidence hint generated by ProtHint (Table S6,ab) as well as the fractions of genes with at least one hint generated by VARUS (Table S7).

When we used protein reference sets with largest sizes, more than 80-85% of annotated genes of *A. thaliana, C. elegans, D. melanogaster, N. crassa* and *D. rerio* harbored protein hints; this percentage was 69% in *S. lycopersicum* (Table S6a). These numbers went down to a range of 40-60% when the sets of reference proteins decreased to their minimal sizes, that were reached when all proteins from the same phylum species were excluded. It was interesting to make comparison of just cited data with the percentage of annotated genes receiving high-confidence protein hints. For the largest reference sets, we observed that percentage of genes with high confidence hint was lower than respective previous figures by just 3-5% in *A. thaliana,* and *D. melanogaster, S. lycopersicum* and *D. rerio;* the drop, however, was by 13% in *N. crassa* and by 24% in *C. elegans* (Table S6b). This large drop for *C. elegans* corresponds to the lowest accuracy of GeneMark-EP+ among all the species considered (Fig. 5b). For the smallest reference sets, proteins from the same phylum excluded, the drop in high-confidence hints coverage was within 10-15% range for all species but *N. crassa* (22%).

The coverage by all protein hints (high-confidence or not) in comparison with coverage by RNA-Seq derived hints (Table S7) was higher by 10-15% for all species but *C. elegans* (lower by 10%) and *D. rerio* (lower by 5.5%). Finally, we saw that annotated genes that did not harbor any hints, either from ProtHint or from VARUS, made a sizable fraction only in *S. lycopersicum* - 24.4% (Table S8)., while in all other species these fractions were rather small: from 5% (*C. elegans)* to 12.3% (*N. crassa*).

In our data, the percentage of annotated genes covered in a given genome by the largest set of protein hints is comparable or higher than the percentage of genes covered by the transcripts derived hints. Also we saw that a vast maj oriy of genes in the six species receive one or another type of external support. The genes that don’t have external support belong to intersections of sets of genes that code for orphan or unique proteins and sets of genes that did not show detectable expression in the experiments measuring gene expression. Still we have to make correction for the fact that RNA-Seq derived hints were not defined for single exon genes even if they were expressed.

### Sources of improvements in gene prediction

Better performance of GeneMark-EP+ in comparison with GeneMark-ES, is expected due to two factors a/ model parameterization on a better validated training set as the training process becomes semi-supervised instead of unsupervised and b/ enforcement of high confidence hints in gene prediction steps. Notably, even when direct corrections are not made (GeneMark-EP mode where factor b/ is absent), for all the species but fungi GeneMark-EP showed improvement over GeneMark-ES. Surprisingly, GeneMark-EP showed only small fluctuations in accuracy when the size of the reference set of protein increased by including more evolutionary close species (Table S3).

The accuracy of GeneMark-EP+ was about the same as the accuracy of GeneMark-EP when the smallest reference set of proteins was used (proteins from species outside the phylum of the species in question). Accuracy of GeneMark-EP+ increases significantly when reference proteins from more evolutionary close species are included while accuracy of GeneMark-EP stays about the same. The only exception was *C. elegans* in which GeneMark-EP gene level accuracy dropped by ~4% for the reference set of species outside the same phylum in comparison with GeneMark-ES (while GeneMark-EP+ shows the accuracy close to the level of GeneMark-ES, Table S3).

These observations suggest that even a relatively small number of anchored introns plays a critical role in parameter estimation in GeneMark-EP. Further increase in the number of anchored introns does not improve parameters of GeneMark-EP. For the case of *C. elegans*, one could argue that the sufficient minimal number of anchored introns was not found when proteins of the reference set were limited to ones from the species outside the *C. elegans* phylum.

To differentiate contributions into GeneMark-EP+ performance, we compared runs that used only high-confidence intron hints with runs that used only high-confidence hints for gene starts and stops (Table S9). This experiment showed that enforceable hints of both kinds contributed equally to overall accuracy improvement. However, these hints contribute unequally into reduction of different types of error. Enforcement of high-confidence intron hints led to higher prediction accuracy of internal exons, while enforcement of high-confidence hints to gene starts and stops led to reduction of errors in initial and terminal exons.

We observed that GeneMark-ES was more likely to generate gene merging than gene splitting errors (Table 4); for instance, comparison of the *A. thaliana* gene predictions and annotation showed 360 split genes and 743 merged genes. Use of GeneMark-EP (with reference proteins outside the same genus) decreased frequency of errors in gene merging (a ~15% decrease in all species) however, it also caused a slight increase in gene splitting (Table 4). Transition to GeneMark-EP+ (the last column in Table 4) reduces gene merging dramatically.

**Table 4:** Numbers of merged and split genes in predictions of GeneMark-ES, -EP and -EP+ with enforcement of (a) only high confidence hints to introns, (b) only high confidence hints to gene starts and stops (c) enforcement of both (a) and (b). All the numbers were obtained for reference sets of target proteins from the species outside of relevant genus.

|  | Genes | ES | EP | EP+<br>Introns<br>(a) | EP+<br>Starts/Stops<br>(b) | EP+<br>Full<br>(c) |
| --- | --- | --- | --- | --- | --- | --- |
| *N. crassa* | Merged | 129 | 89 | 92 | 64 | **74** |
|  | Split | 83 | 96 | 89 | 106 | **91** |
| *C. elegans* | Merged | 1120 | 1076 | 1090 | 1019 | **1029** |
|  | Split | 588 | 725 | 614 | 731 | **622** |
| *A. thaliana* | Merged | 743 | 634 | 629 | 215 | **251** |
|  | Split | 360 | 385 | 242 | 478 | **277** |
| *D. melanogaster* | Merged | 544 | 464 | 462 | 311 | **313** |
|  | Split | 285 | 297 | 204 | 324 | **221** |
| *S. lycopersicum* | Merged | 2304 | 1871 | 1793 | 1165 | **1192** |
|  | Split | 1550 | 1644 | 1139 | 1962 | **1252** |
| *D. rerio* | Merged | 1921 | 1415 | 1351 | 883 | **884** |
|  | Split | 2553 | 2976 | 2018 | 3058 | **2010** |

Enforcement of only high-confidence intron hints in GeneMark-EP+ reduced the number of split genes (by enforcing introns in place of incorrectly predicted intergenic regions). Still these hints have little or no effect on the gene merging (Table 4). The most significant effect on gene splitting was observed for *D. rerio* - 2010 split genes in the -EP+ mode compared to 2976 in the -EP mode.

Enforcement of high confidence hints to gene starts and stops significantly reduced numbers of merged genes and caused a slight increase in numbers of split genes. For instance, the number of merged genes dropped by ~500 in *A. thaliana* between GeneMark-ES and GeneMark-EP+, a ~66% improvement; about 50% improvement was observed for the other species in our tests except *C. elegans*. All over, GeneMark-EP+ (Table 4, last column) achieved significant reduction in numbers of both merged and split genes in comparison with GeneMark-ES and GeneMark-EP.

### Comparison of GeneMark-EP+ predictions with genome annotations defined by the APPRIS database

We compared GeneMark-EP+ gene predictions with annotations of major protein isoforms in *C. elegans*, *D. melanogaster, and D. rerio* genomes defined by the APPRIS database (24). This test did show (Fig. S6) an increase in exon level sensitivity (by ~4% for *C. elegans and D. rerio,* by ~7% for *D. melanogaster)* and a decrease in exon level specificity (by ~1.5% for *C. elegans,* by 3% for *D. melanogaster* and by ~8% for *D. rerio)* in comparison our previous assessment results using genome annotations made by respective genomic communities (Table 1). The decrease in Sp could be expected since the APPRIS annotation contains smaller number of exons. The increase in Sn is a positive news indicating that GeneMark-EP+ when making prediction of a single isoform per locus is likely to predict genes for major protein isoforms. At gene level (Fig. S7), both Sn and Sp were reduced slightly in *C. elegans* and *D. rerio,* and by 5% in *D. melanogaster.* To correctly interpret this result, we have to remind the definition of gene level accuracy - a gene is counted as correctly predicted if the prediction matches all exons in at least one alternative transcript. Thus, a gene is considered to be predicted correctly if just one of the isoforms (major or not) is correctly predicted (Fig. 5, Table S3). This is a rather liberal way of computing an Sn value on gene level.

### Assessment of accuracy of ProtHint

The main role of ProtHint is generation of a list of co-ordinates as well as confidence scores of potential borders between coding and non-coding regions in a novel genome. Specific thresholds on confidence scores are define to select subsets of hints (e.g. high-confidence set). The GeneMark-EP training procedure can tolerate a high number of false positive intron hints since only a subset, the anchored introns, are used in training. It is important that the set of *all mapped hints* would have high Sn with respect to true gene elements while the Sp level could be lower. On the other hand, the *high-confidence hints*—those utilized in initial GeneMark-EP+ parameter estimation as well as in the hints enforcement—have to have high Sp, as these hints are directly enforced in predictions.

### How large are fractions of correct hints among hints generated by ProtHint?

When the set of reference proteins had the maximum size (all proteins in a relevant OrthoDB division except ones from the same species) the set of intron hints generated by ProtHint had Sn > 75% for exact introns and Sn ~70% for gene starts and stops (Tables 5, S10). The value of Sn was dropping down steadily as evolutionary distance to reference proteins was increasing. Particularly, when the proteins from species of the same *order* were excluded, Sn was, on average, ~65% for intron hints and ~40% for gene start and stop hints.

**Table 5:**
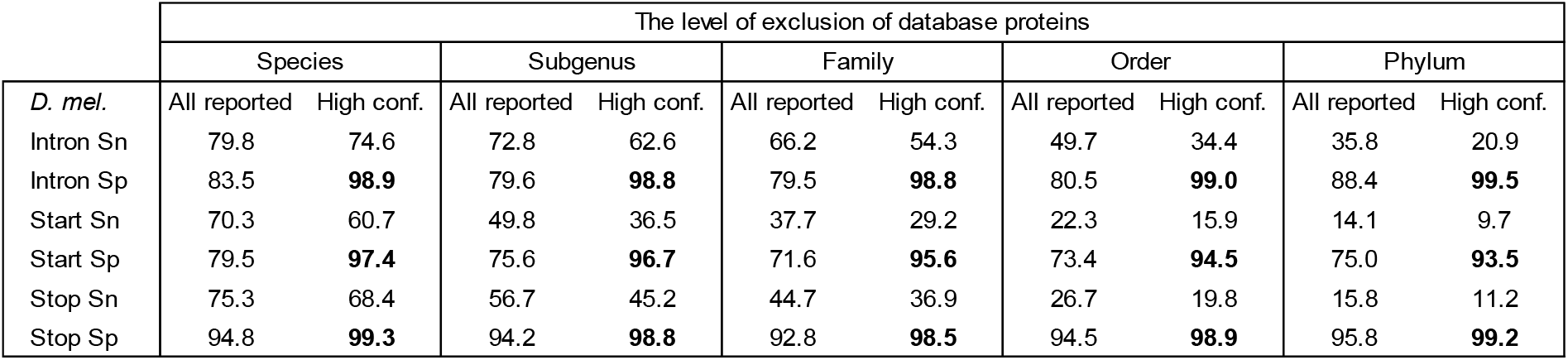
Accuracy of ProtHint for the *D. melanogaster* genome: sensitivity and specificity of hints to introns, start and stop codons. The results are shown for all reported hints or just high-confidence hints. The Sn and Sp values are computed based on genome annotation of a full complement of introns, gene starts and stops, including alternative isoforms. Results for all tested species are shown in Table S7.

The largest reduction in the volume of the protein reference set – exclusion of proteins from the same phylum - decreased Sn of all reported intron hints down to ~40% on average (Table S10). Here the largest Sn value (the fraction of correct intron hints) was observed for *N. crassa* (60%), the lowest one - for *C. elegans* (26%). At the same time, the value of Sn of gene start and stop hints generated from the smallest reference set of proteins varied significantly between the species, from 8% for *C. elegans* to 30% for *N. crassa* (Table S10).

### How reliable are ‘high confidence’ hints generated by ProtHint?

The sets of high-confidence hints were observed to have high specificity, averaging over 95% (5% of false positives) over the six species. This level remained high even for the smallest sets of reference proteins, proteins from the species outside the phylum of interest (Tables 5, S10). In case of *C. elegans,* along with high Sp, we observed low Sn value of high-confidence hints (for all the reference protein sets – larger or smaller) which is explained by the presence of just a few species with sequenced genomes in the *C. elegans* phylum (Table 2). For all other species, adecrease in Sn upon transition from *all mapped* to *high-confidence* hints was small in comparison with the simultaneous increase in Sp.

Distributions of (IMC, IBA) vectors representing intron hints generated for *N. crassa* (both false and true as compared with annotation) are shown in Fig. 4a (for the genus-excluded protein reference set). The Sp-Sn curves are generated for sets of introns hints obtained by filtering with IMC and IBA thresholds (Fig. 4b).

The distribution of the score vectors (Fig. 4a) as well as the behavior of Sp-Sn curves (Fig. 4b) depend on selection of the set of reference proteins (genus or order or phylum excluded, Figure S2, left and middle panels). A choice of IBA threshold selecting high-confidence intron hints affects accuracy of GeneMark-EP+. We assessed the extent of this effect for *A. thaliana*, *N. crassa*, and *S. lycopersicum* (Figure S2, right panels). It was shown that the best average prediction accuracy was achieved with IBA threshold set to 0.25. Similar analysis produced necessary thresholds for high-confidence hints to gene starts and stops.

### More intron hints are generated in regions encoding conserved protein domains

We found that about 50% of the whole set of introns annotated in the APPRIS set of principal isoforms is located within conserved protein domains (Table S11).

In *D. melanogaster,* high-confidence intron hints generated by ProtHint from the ‘species-excluded’ reference set of proteins fell into regions coding for conserved domains in 55.9% of cases (Table 6). This fraction increased significantly as more proteins were excluded from the reference set (e.g. proteins from species outside of the *D. melanogaster* genus). This fraction reached 84.6% when only proteins originated from species outside the *D. melanogaster* phylum were considered (Table 6). Similar trends were observed for *C. elegans* and *D. rerio* (Table S12). In the set of *all reported* intron hints, the fraction of introns mapped to regions coding for conserved domains was lower than in the set of high-confidence intron hints (Table S12), however, the proportion of introns mapped into conserved domain regions also increased upon removing proteins from closely and moderately closely related species.

**Table 6:** For D. *melanogaster* genome we show fractions of high-confidence (HC) intron hints mapped into regions coding for conserved protein domains. The results are provided for sets of reference proteins with different sizes and evolutionary distance to *D. melanogaster.* Out of 41,010 introns in the APPRIS *D. melanogaster* genome annotation, 21,562 (52.6%) are located in regions encoding conserved protein domains.

| Exclusion level | High-confidence introns matching APPRIS introns |  |  |
| --- | --- | --- | --- |
|  | All HC introns | HC introns that fell into domains |  |
| Species | 33,894 | 18,934 | (55.9%) |
| Subgenus | 28,437 | 17,475 | (61.5%) |
| Family | 24,670 | 16,057 | (65.1%) |
| Order | 15,829 | 11,984 | (75.7%) |
| Phylum | 9,719 | 8,222 | (84.6%) |

The same type fractions computed for ‘high-confidence’ and for ‘all reported’ intron hints were almost identical for the species between *D. melanogaster* and *D. rerio.* (Table S12). Still, for *C. elegans,* however, the fraction of true high-confidence introns was lower (Table S12) apparently due to having fewer target proteins from close relatives in the protein database (a factor significantly affecting IMC score).

## Discussion

The main reason to develop GeneMark-EP, EP+ was a clear need to leverage abundant protein sequence data available in public databases for improving accuracy of automatic gene prediction. It was well expected that iterative *ab initio* parameterization of statistical models (as done in GeneMark-ES) would become more precise, especially for large genomes, if we find an efficient method to add data on protein footprints into training and prediction steps. This project has grown into development of a whole GeneMark-EP pipeline, with GeneMark-EP+ mode that directly integrates most confident evidence into predicted exon-intron structures. In this respect, the new pipeline features a new method, ProtHint, developed to find multiple proteins homologous to a gene initially predicted in a genomic locus and then to derive reliable hints to the true gene exon-intron structure by constructing and processing multiple protein footprints. GeneMark-EP, EP+ should become a universal extension of GeneMark-ES, as its application to a novel eukaryotic genome will be facilitated by use of a vast volume of protein sequences.

Another earlier developed method, GeneMark-ET (2), extended GeneMark-ES to use external evidence generated from transcriptome sequence data, when it is available along with a newly assembled genome.

Existing methods, such as GenomeThreader (7), rely on mapping proteins from closely related species as well as mapping gene elements from aligned genomic sequence of the close species to produce predicted exon-intron structures. However, its prediction accuracy is dropping fast with increase of evolutionary distance between species (6).

Use of multiple homologous proteins proved to be important for keeping decent accuracy of prediction with increase of evolutionary distance between species with known genomes and the species of interest. Particularly, due to corroboration of footprints originated from multiple homologous proteins, we observed enrichment of high-confidence introns in regions coding for conserved domains (Table 6).

Use of anchored elements of gene structure was important for integration of signals originated from different sources (sites predicted from genomic sequence alone and sites identified by protein footprints). The logic of selection of anchored elements enabled filtering out ‘one-sided’ noises present in one or another source. Use of partial protein footprints, when a target protein mapping could contribute less than full exon-intron structure was another important feature of the new method. Partial footprints were useful for improving training sets; they also added confident corrections at gene prediction steps (Fig. S8).

Use of anchored elements was most beneficial for large genomes (*S. lycopersicum* and *D. rerio)* where GeneMark-ES alone generated an elevated rate of random false positive errors within long intergenic regions.

Mapping of N- and C-terminals of target proteins allowed for better discrimination between introns and intergenic regions than it could be done by an *ab initio* algorithm. This improvement led to significant reduction of errors in gene merging (when intergenic regions were predicted as introns) though reduction in error rate of gene splitting (when introns were predicted as intergenic regions) was smaller (Table 4).

The most significant improvement in comparison with GeneMark-ES, observed in all species but fungi, *N. crassa,* occurred when GeneMark-EP+ used the largest possible set of reference proteins (Figs. 5, S5). For *N. crassa,* use of protein evidence never led to noticeable improvement over GeneMark-ES which high accuracy for fungal genomes was demonstrated earlier as well (21). We assume that relative drop in GeneMark-EP, -EP+ performance for *C. elegans* in comparison with *Arabidopsis* and *Drosophila* was related to a lower number of reference proteins within the *C. elegans* phylum. In tomato and fish genomes that have longer on average intergenic regions than other species we saw low exon level specificity (~55-60%) related to elevated false positive prediction of protein-coding genes in long intergenic regions (Fig. S5). Gene level accuracy for *D. rerio,* ~30% Sn and ~12% Sp, for any set of reference proteins beyond the *D. rerio* genus, was difficult to improve. Notably, the genes in fish genome have a rather large, 8.2, average number of introns per gene. Under independence of errors assumption, a gene with a large number of introns would be improbable targets for accurate prediction. Even though, the independence assumption does not hold in presence of external evidence, the gene error rate increase with the increase in number of introns (data not shown).

Annotation of genes encoding principal protein isoforms is available for *D. melanogaster*, *C. elegans* and *D. rerio* in the APPRIS database (24). GeneMark-EP+ comparison with the APPRIS annotation shows better Sn than in comparison with annotations containing all possible isoforms.

A question could be raised, how do pseudogenes affect training of GeneMark-EP, EP+. This question is difficult to address in a general setting. Still, since pseudogenes could have different age, let consider just groups of ‘young’ and ‘old’. Young pseudogenes with one or two mutations that make them dysfunctional still have all the sequence patterns that could be used in training. Old pseudogenes that accumulated many mutations would harm statistical models if included in training. We argue that old pseudogenes will not be predicted by GeneMark-ES in the course of self-training and therefore they have little or no chance to be included in a training set of anchored elements. On the other hand, elements of young pseudogenes could be identified by GeneMark-ES while the frameshifted exons from spliced alignments will be detected and scored unfavorably by ProtHint. Therefore, the young pseudogenes could contribute to parameter training as their ‘intact’ parts will appear in both training and prediction. Addressing full complexity of this issue goes beyond the scope of this project, therefore, currently, GeneMark-EP, EP+ do not collect information on frameshifts and potential pseudogenes.

Interestingly, the second run of full GeneMark-EP+ (when we took as seed genes the results of gene predictions made in the first full run) had a small but positive effect on the final gene prediction accuracy. This additional run is recommended if an increase in run-time is not a concern.

Running GeneMark-EP, EP+ requires a protein database as well as tools searching for target proteins and for protein splice alignments. We used OrthoDB as a database of reference proteins, DIAMOND (25) for the database search for proteins (targets) homologous to the seed proteins and Spaln (9) for splice alignment of target proteins to genome. To accelerate the pipeline run we limited the DIAMOND output by25 target proteins per seed protein (Fig. S9); choice of Spaln was also practical from the standpoint of run-time reduction. Additionally, we verified that use of GeneMark-ES for generating seeds was a faster and more efficient method in comparison with the six frame translation with Procompart and ProSplign tools (8).

This discussion section would be incomplete if we do not mention limitations of the new method. GeneMark-EP does not support multiple models needed for genomes with heterogeneous nucleotide composition, like genomes of mammals and some plants (grasses, e.g. rice). While the current version of GeneMark-EP, -EP+ would outperform GeneMark-ES when running on such genomes, the overall accuracy could be significantly improved with more accurate modeling of genome heterogeneity.

We realize that use of taxonomic divisions for selecting or out-selecting of reference proteins is just the first step in accurate modeling of real-life distributions of evolutionary distances to database orthologues for genes and proteins existing in a novel species. Arguably, there is room for improvement of both intron and gene start/stop hints when modeling of sets of reference proteins is done based on evolutionary distance measures. Similarly, one would expect that such modeling would lead to improving in selecting thresholds for introns and sites mapping.

Another limitation of the current method is the search for a single optimal genomic sequence parse that leads to prediction of a single gene and a single protein isoform in each locus. Importance of genes with alternative splicing has been debated recently, as the evidence was accumulated that alternative splicing mainly operates with UTR regions rather than with translated regions of pre-mRNA. Moreover, the claims were made that when a translated region could be alternatively spliced then only one among the protein isoforms, the major one, is expressed in the largest number of tissues (24). If gene prediction by GeneMark-EP, -EP+ is viewed as prediction of the major isoform, then the result should be naturally assessed in comparison with annotation of the major isoforms. Such comparison, done for *C. elegans, D. melanogaster, and D. rerio*, used annotation provided by the APPRIS database, and showed improved sensitivity in predicting genes of major protein isoforms. Nonetheless, general tools able to predict all alternative isoforms are of significant interest for community. When external information representing alternative isoforms is provided at RNA level, an earlier developed pipeline, BRAKER1 (32), uses GeneMark-ET and AUGUSTUS to make predictions of alternative isoforms. A new pipeline, BRAKER2 (paper in preparation) combines GeneMark-EP, -EP+ with AUGUSTUS to identify a set of alternative protein isoforms when alternative variants of cross-species proteins are given among references. A new tool, GeneMark-ETP, will combine into gene prediction protein and transcript data (paper in preparation).

## Availability

Full GeneMark-EP, -EP+ package, including ProtHint, is available at http://topaz.gatech.edu/GeneMark/ license_download.cgi. Software is compiled for Linux and Mac OS operating systems. All scripts and data used to generate figures and tables in this manuscript are available at https://github.com/gatech-genemark/GeneMark-EP-ProtHint-exp. To give an example, the overall runtime of ProtHint and GeneMark-EP in EP+ mode on the *D. melanogaster* genome (having ~14,000 genes in 134 MB sequence) with target proteins selected from species outside *Drosophilidae* family was ~5 hours on 8CPU/8GB RAM machine. In our experiments the run time grew linearly with respect to both genome length and number of genes.

## Supplementary Materials

Supplementary Materials are available at the NAR G&B web site.

