## Supplemental Materials for "GeneMark-EP and -EP+: eukaryotic gene prediction with self-training in the space of genes and proteins"

#### Additional Changes Made in GeneMark-EP, -EP+ in Comparison with Published Versions of GeneMark-ES, -ET

In the published versions of GeneMark-ES, -ET algorithms an intergenic region length distribution was modeled by a uniform distribution. Currently, non-parametric estimation of probability density function of the length distribution is used in GeneMark-ES, -ET, -EP and -EP+ for all species but fungi. The non-parametric estimation is applied in the last (third) round of self-training iterations (see flowchart in Fig. 1). Uniformly distributed pseudocounts are added to smooth the distributions. In GeneMark-EP, -EP+, only those intergenic regions that are situated between genes with anchored introns are used as data for the non-parametric estimation of the length distribution.

Default minimum length of genes predicted by GeneMark-ES, -ET was set to 300 nt. In GeneMark-EP, EP+, shorter genes will appear in the list of predicted genes if they are supported by at least one High-Confidence hint.

GeneMark-ES/ET/EP, -EP+ has a model for non-canonical GC-AG introns.

#### Accuracy Assessment on Exon Level

Exon level (Fig. S5) accuracy dependence on changes in size of sets of reference proteins followed the same trends as the gene level accuracy dependence (Fig. 5, description in the main text).

**Fungal genomes**: *N. crassa*. In comparison with GeneMark-ES we observed small improvements in GeneMark-EP+ (by ∼2%) when the hints originated from the proteins of species outside *N. crassa* genus and order (Fig. S5a). No difference between -ES and -EP+ was observed when the hints were coming from proteins outside of the *N. crassa* phylum.

**Compact eukaryotic genomes**: *C. elegans*, *A. thaliana*, and *D. melanogaster*. GeneMark-ES was quite accurate within this group of genomes, still the prediction accuracy of GeneMark-EP+ was higher. The improvement was most pronounced for *A. thaliana* (Fig. S5c) and *D. melanogaster* (Fig. S5d). GeneMark-EP+ with hints from proteins from the relevant genus and beyond improved over GeneMark-ES by 5%-10% in both Sn and Sp. This improvement was reduced when evolutionary distance to target proteins increased. However, even for more distant target proteins situated outside the relevant phylum, we saw increase in specificity by

2%-4%. For *C. elegans* (Fig S5b) the accuracy of GeneMark-EP+ improved slightly over -ES when target proteins from inside the same genus were admitted but remained almost the same when target proteins were selected from species outside the *C. elegans* genus or phylum.

**Large eukaryotic genomes**: *S. lycopersicum* and *D. rerio.* GeneMark-ES was less accurate for large genomes than for compact genomes. When proteins from the species inside the same genus could be used as targets for hints generation, GeneMark-EP+ showed significant increase in performance (Figs. S5ef) with Sn ~75% comparable to the Sn values reached for the compact genomes. The Sp value was improved to 55%-60%. Still, it was much lower than the average Sp observed for compact genomes. When target proteins were selected from species outside the relevant genus, improvement of GeneMark-EP+ over GeneMark-ES was by ~10% in Sn and Sp (Figs. S5ef). This improvement remained high even when more remote target proteins were used for hints generation, i.e. from species outside the same order or outside the same phylum.

### Details of the ProtHint Design

#### Scoring of Intron hints

##### Alignment of Entire Exon (AEE)

The AEE scores are computed for all exons adjacent to introns mapped by spliced alignments. An AEE score is defined for a whole exon appeared in alignment constructed by Spaln for a target protein. The alignment score is computed with BLOSUM62 parameters and linear gap penalty $-4$. An AEE score is not normalized by the exon length, therefore, exons with low scores are either too short or they are long and poorly aligned. At the initial step of the algorithm we keep introns bordered by exons with high AEE scores.

If an exon has a frameshift (or internal stop codon), the Spaln generated spliced alignment detects the change of the reading frame (or stop codon). In this case we split the exon into two parts and score the parts separately. These two scores, if high enough, could be used for IBA scoring of corresponding adjacent introns.

##### An IBA Score for an Exon with a Frameshift

If a frameshift (indel) is detected by the spliced alignment algorithm, we modify the protein alignment downstream from the frameshift point (for downstream exon) or upstream from the frameshift point (for upstream exon) by replacing each translated codon with a gap. Each such artificial gap adds penalty $-4$.

##### Comparison between Intron Border Alignment (IBA) and Intron Mapping Coverage (IMC) Scores

Comparison between IBA and IMC scores showed that a high value of IMC is a better indicator of high intron specificity than high value of IBA (Fig. 4). A combination of these two scores allowed us to relax the IMC threshold and to get a larger set of High-Confidence introns.

A direct comparison of Sp-Sn curves (Fig. 4) is not entirely fair for the following reason. All introns are filtered with IBA $\geq 0.1$ and AEE $\geq 25$ prior to computing IMC (Fig. S1). This removes a significant number of false predictions. The IMC score is computed from a set already filtered with IBA score.

#### Computation of Start Overlaps

Precise definition of a gene start overlap by a target protein footprint is as follows. Start S is considered to be overlapped by a target protein P if an exon E in P overlaps S upon spliced alignment. Still, to be counted as overlapping, exon E needs to satisfy these criteria: (i) AEE score of E has to be $\geq 25$. (ii) The spliced alignment of protein P must contain a mapped start codon or an acceptor site (within the set of all reported starts/introns) which coincides with the exon start. In other words, the start of an exon must define either a start codon or an acceptor splice site. See Figure S3 for illustration.

#### Threshold Selection for Ensuring High Specificity of High-Confidence Hints

Errors in spliced alignment which create falsely mapped introns do not significantly influence GeneMark-EP training. Training with use of all introns from ProtHint vs training with use only true ProtHint introns generates almost the same GeneMark-EP accuracy. GeneMark-EP+ improves over GeneMark-EP mostly due to influence of hints on prediction steps and in much less extent due to improvement of training. High specificity of the high confidence hints is critical for the hints to work. Therefore, a significant effort was made to develop high confidence selection criteria, notably:

- The coverage scores are somewhat similar to intron coverage score for RNA-Seq reads intron mapping in GeneMark-ET (1). The IMC threshold "$\geq 4$" was tested for proteins from two databases: EggNOG and OrthoDB. In both cases, the cited threshold was leading to similar results across various species tested.
- Scores IBA and BAQ used in high confidence hint selection characterize the quality of spliced alignment near the co-ordinates of a candidate hint. We selected a linear (triangle) kernel, which gives higher weight to alignment positions close to coding region boundaries. We tested several other kernels (box, parabolic, triweight), however, the linear one was generating consistently best results for windows with different sizes. Comparison between results of application of a linear and box kernel is shown in Figure S10. The linear kernel was also most robust with respect to changing window sizes. Window sizes 5, 10, 15, and 20 were tested and 10 was selected as consistently best performing across species tested.
- We tested several methods for filtering introns as well as alternative formulas for computing intron borders alignment score (IBA). Longer alignments of individual exons did not produce better intron prediction quality (Fig. S1). The IBA score constructed as an arithmetic mean of upstream and downstream scores $S_d$ and $S_u$ was less accurate than a score using geometric mean of $S_d$ and $S_u$.
- For start codon hints, removing starts overlapped by exons from some proteins alignments was critical for ensuring high specificity (Fig. S3, Tables 3, S2)

#### Invariance of the Spliced Alignment with Respect to Alignment Tools

ProtHint also supports use of ProSplign (3) as an alternative to generating spliced alignment with Spaln (2). We observed that accuracy of hints generated by ProSplign as well as accuracy of subsequent GeneMark-EP+ gene predictions did not differ significantly from the results obtained with Spaln. Since Spaln is significantly faster, it is used by default. ProtHint also supports an alternative to DIAMOND (4), a more sensitive but slower BLASTp (5). We have not observed significant difference in ProtHint accuracy when either DIAMOND or BLASTp was used. Since DIAMOND is several orders of magnitude faster than BLASTp, ProtHint uses DIAMOND by default.

#### Differences between the Usage of ProSplign and Spaln

ProSplign has a built-in filtering procedure, therefore the filtering steps described in Methods can be skipped and all hints mapped by ProSplign will be used directly. Still, the procedure of scoring and selection of high confidence hints remains the same.

Slow speed of ProSplign hampers its use. ProSplign does not use heuristics to speed-up dynamic programming based alignment algorithm, therefore it is 10-100x slower than Spaln, depending on the length of the genome locus and the length of protein being aligned. To run ProSplign at a reasonable time the "ProSplign mode" works as follows. ProtHint first runs Spaln to generate a set of hints. For each hint mapped by Spaln, top ten supporting proteins are selected and aligned with ProSplign. This selection reduces the number of target proteins to be aligned by ProSplign by an order of magnitude.

We observed that the raw set of hints mapped by ProSplign was generally less sensitive and more specific than hints produced by Spaln, due to ProSplign's internal filtering procedure. However, the set of high confidence hints was almost the same for both tools, meaning that our scoring system was insensitive to a choice of a spliced alignment engine. Consequently, the results of GeneMark-EP, EP+ did not significantly change when either Spaln or ProSplign based alignments were used. Currently, Spaln is used as a default option in ProtHint due to its higher speed.

##### Use of a Custom Protein Database

A custom protein database could be used as an alternative to OrthoDB (6). A special attention should be paid to construction of such database, as presence of identical proteins (for example proteins from subspecies of the same species) can lead to artificially inflated coverage as well as increase in execution time.

### Supplementary Figures

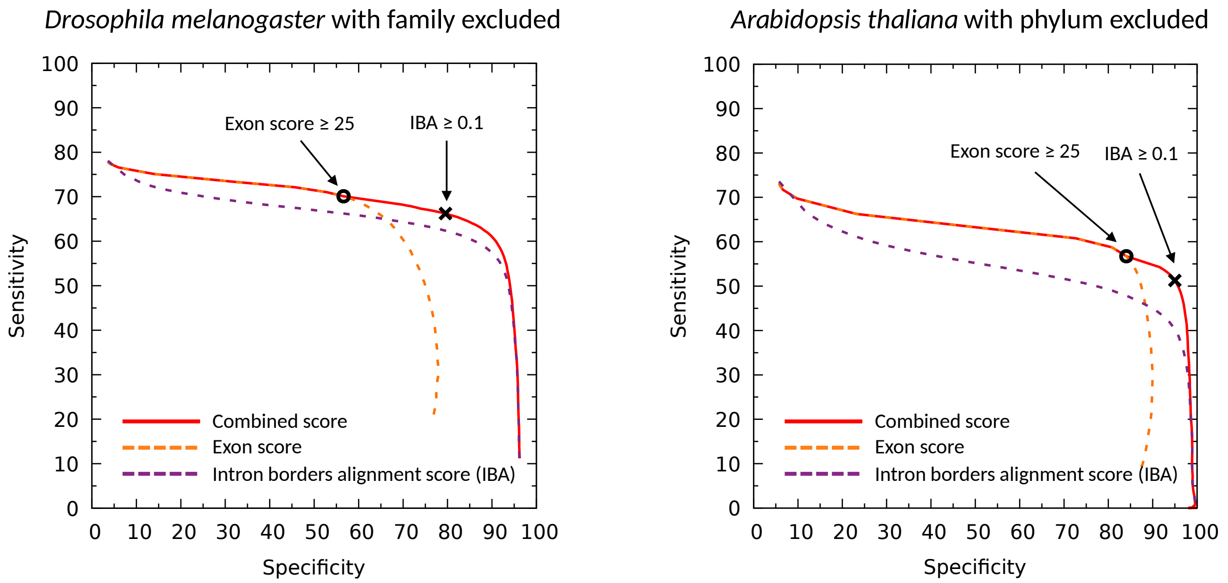


Figure S1: ProtHint intron Sp-Sn curves built upon filtering sets of mapped introns by exon AEE scores (dashed orange) and intron borders alignment score (IBA, dashed purple). The combined curve (red) is generated by, first, selecting out all introns with AEE scores above the threshold changing from 0 to 25; next, all the selected introns are checked for having IBA scores above the threshold changing from 0 to 0.1 and up to 1.0. The position of the black cross in the combined curve represents IBA score $\geq 0.1$ and AEE score $\geq 25$.

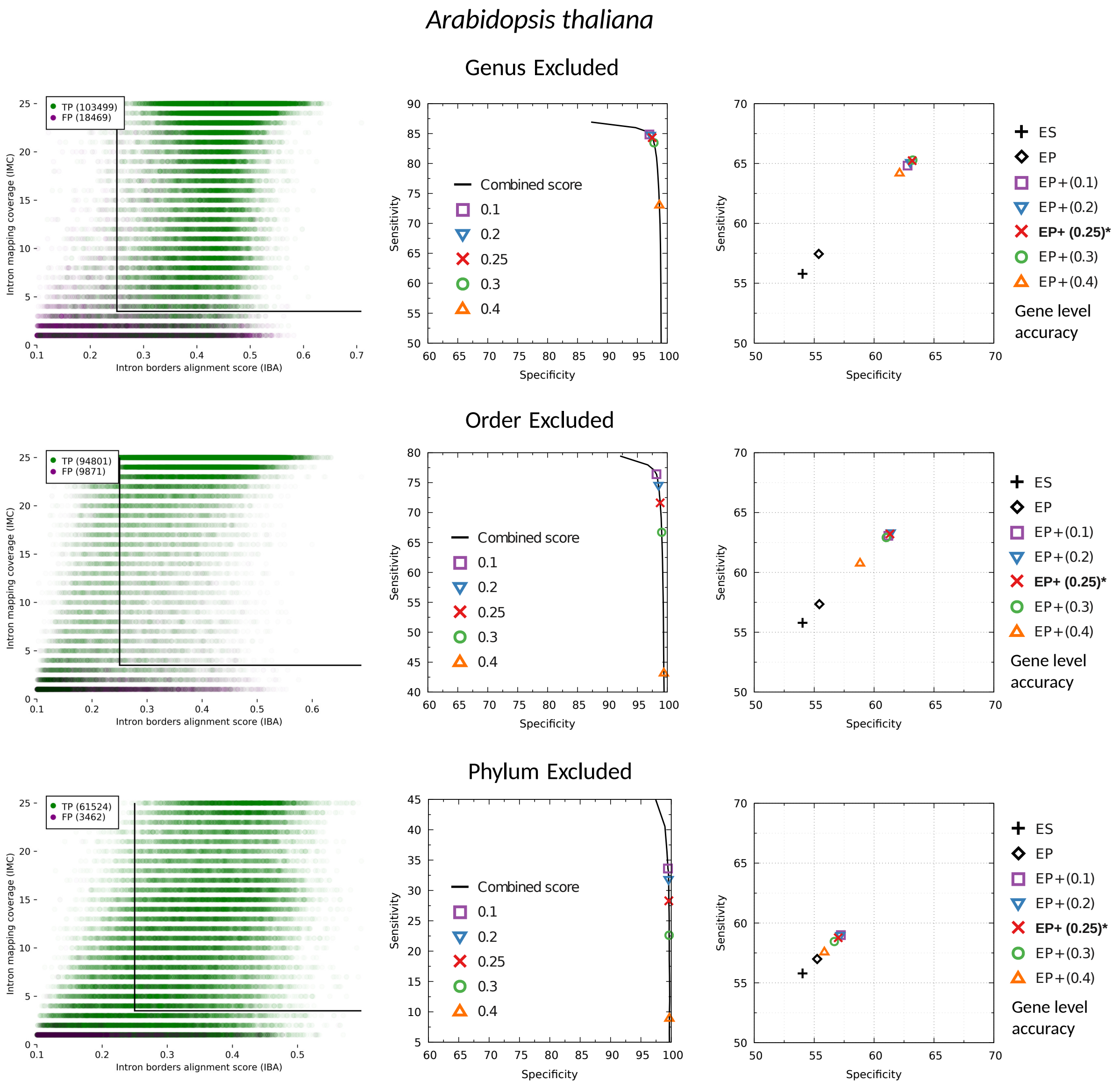


Figure S2a: Effect of IBA threshold on accuracy of high-confidence hints and GeneMark-EP+ for *A. thaliana*.

Left side graphs show distributions of the score vectors of true positive (green) and false positive (purple) introns (mapped and scored by ProtHint), the vectors components are intron borders alignment (IBA) and intron mapping coverage (IMC) scores. The black lines represent cutoffs at IMC = 4 and IBA = 0.25. Total numbers of false and true positives are shown in the upper left corners.

Middle graphs display ProtHint' Sp-Sn curves. The curves are generated by first, selecting out all introns below changing IMC threshold from 0 to 4 and then selecting out all the introns with IBA score from 0 to 0.25 and up to 1.0. The Sp-Sn values for various IBA cutoffs (0.1, 0.2, 0.25, 0.3, 0.4) are shown at the curves. The curves illustrate procedure of selecting introns mapped with high confidence.

Right side graphs display how gene level prediction accuracy of GeneMark-EP+ depends on IBA score cutoffs used to select sets of high confidence introns. Sp and Sn of GeneMark-EP, i.e. without high confidence intron enforcement, as well as for GeneMark-ES, are shown as well.

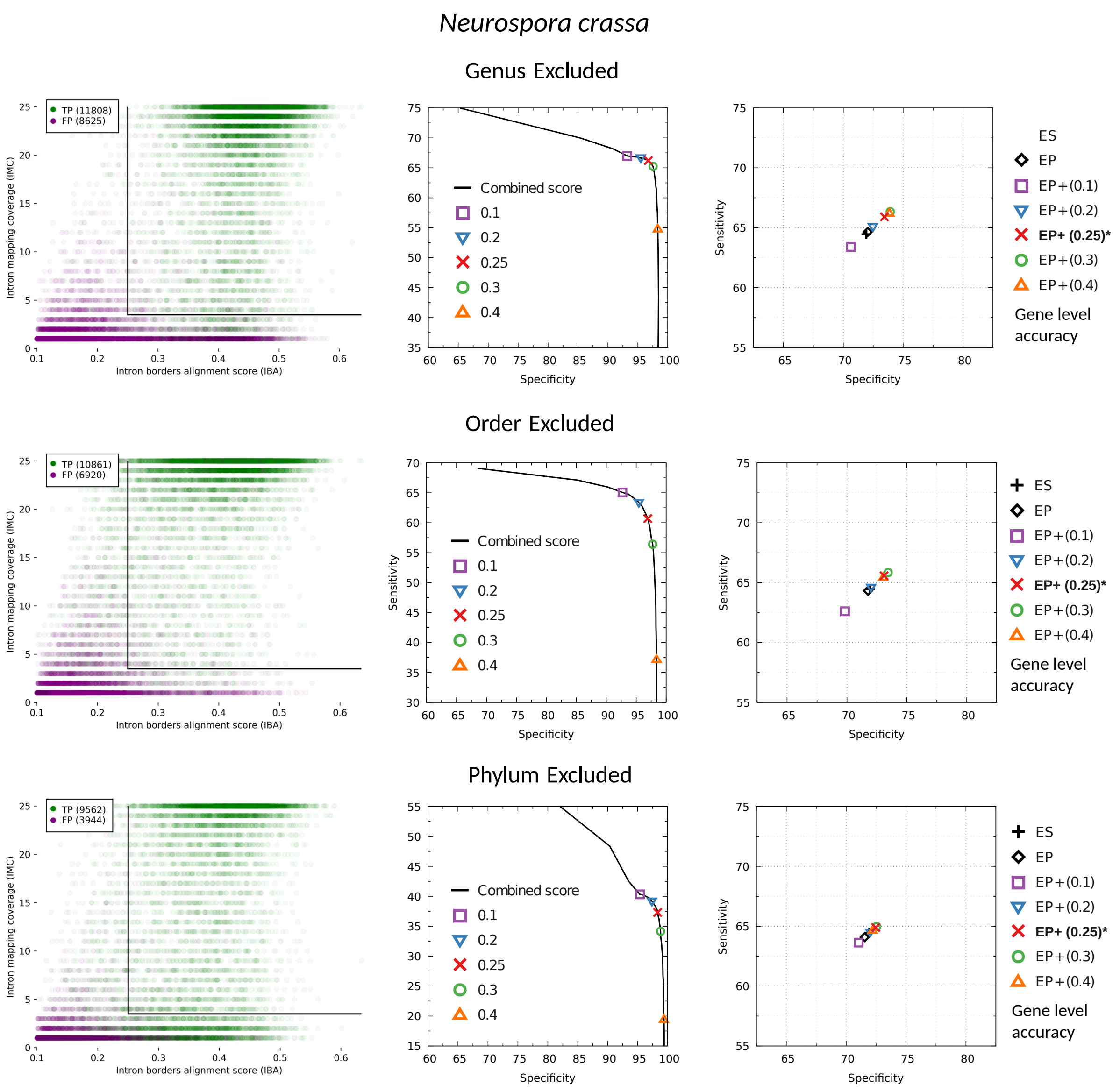


Figure S2b: Effect of IBA threshold on the accuracy of high-confidence hints and GeneMark-EP+ for *N. crassa*. For more details see the legend to Figure S2a.

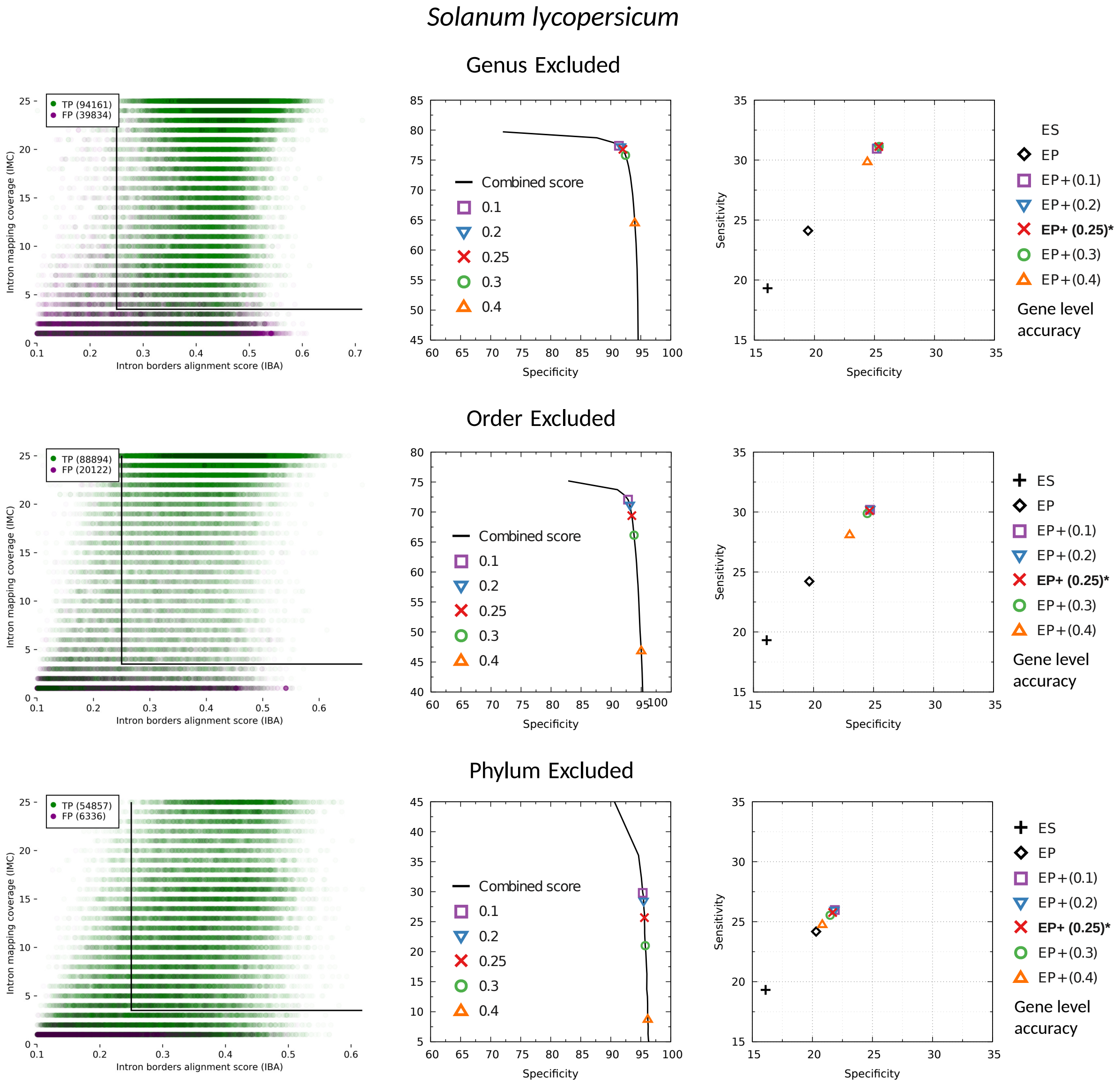


Figure S2c: Effect of IBA threshold on the accuracy of high-confidence hints and GeneMark-EP+ for *S. lycopersicum*. For more details see the legend to Figure S2a.

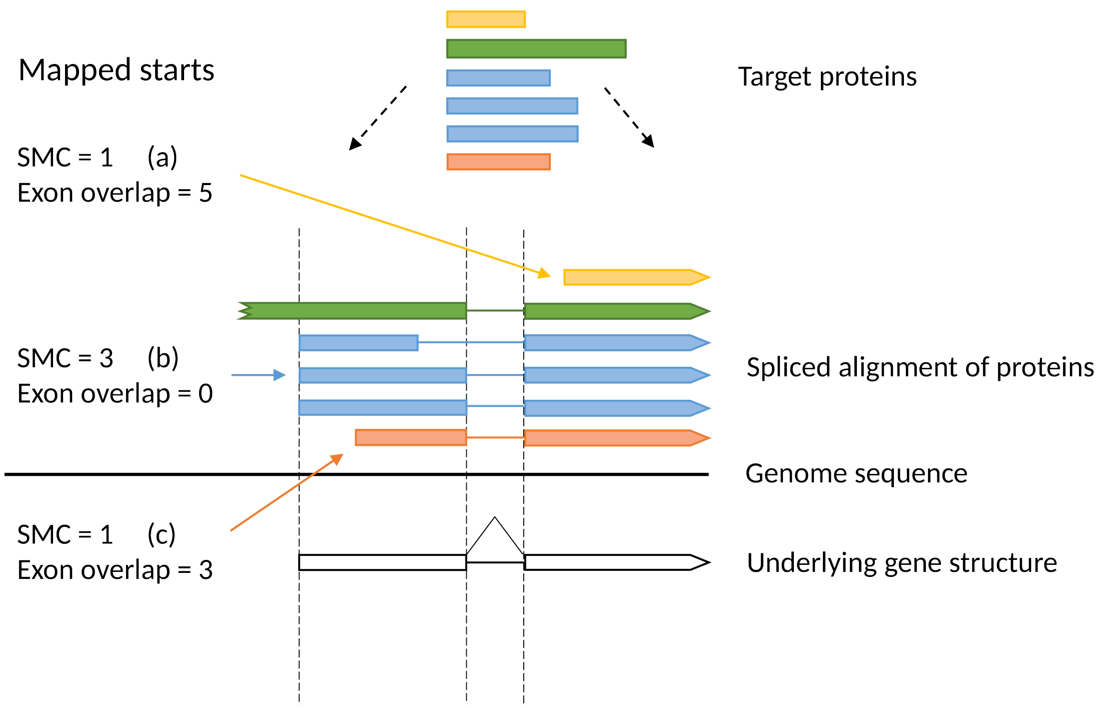


Figure S3: Gene start mapping coverage (SMC) scores and counts of exon overlaps. Start (a) is overlapped by five exons which indicate a presence of an upstream intron. Start (b) is overlapped by one exon (green) but this exon's upstream start does not coincide with an end of an intron or a start codon mapped by ProtHint, therefore it does not contribute to the exon overlap and the exon overlap of start (b) is at zero and start (b) is selected. Start (c) is overlapped by three exons which define an upstream start, green exon is again not counted, thus, start (c) is ignored.

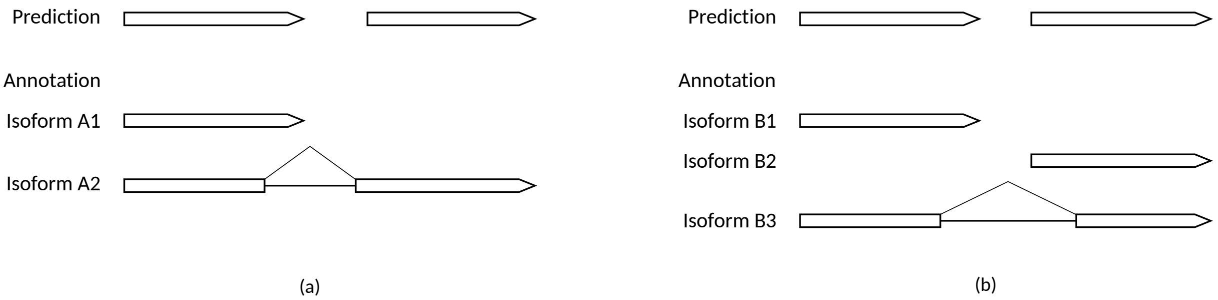


Figure S4: Gene splitting events caused by alternative isoforms that include other isoforms as their components. We remove such cases from the test set for gene splitting assessment. (a) Isoform A1 is correctly predicted. As a result, full isoform A2 cannot be predicted at the same time and it is split. (b) Algorithm makes correct predictions of isoforms B1 and B2. If isoform B3 is considered as annotation, it is split in prediction. For genes with annotated multiple alternative isoforms we use the longest one as a representative

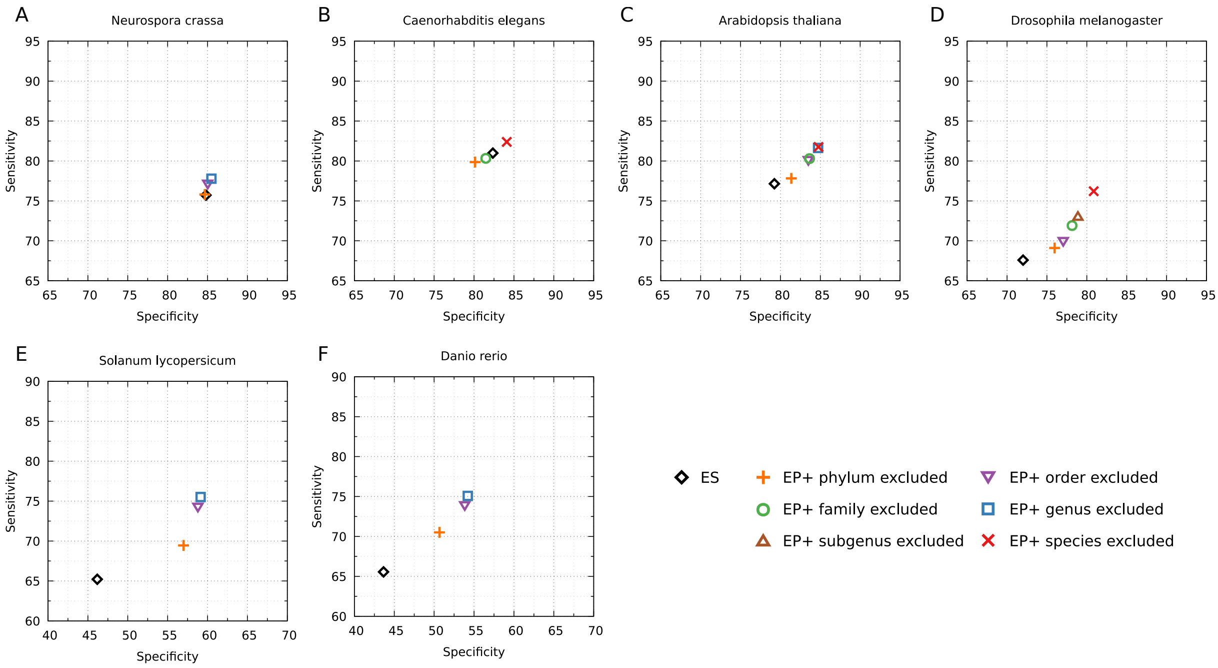


Figure S5: Comparison of GeneMark-ES and GeneMark-EP+ accuracy on exon level. Accuracy of GeneMark-EP+ is shown for cases when ProtHint works with different in size sets of reference OrthoDB proteins: from the largest (only the same species excluded) to the smallest (the whole same phylum excluded). Exon level Sn and Sp are defined with respect to a full complement of annotated exons, including alternative types.

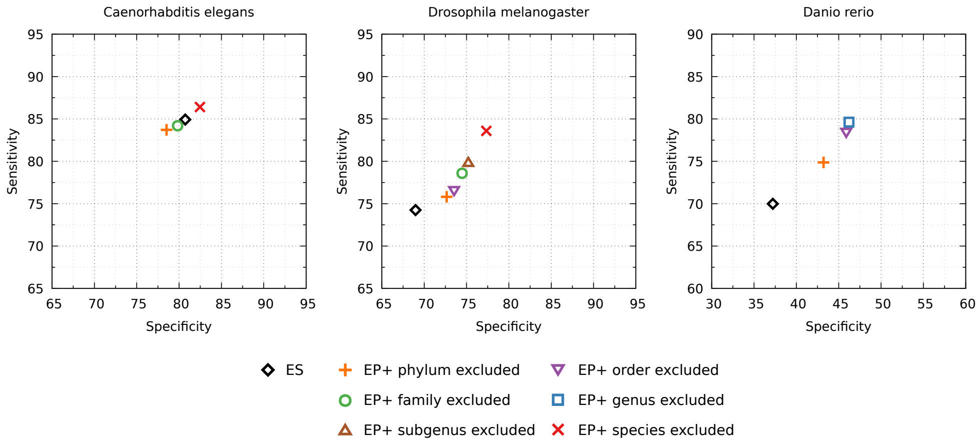


Figure S6: Same comparison as in Figure S5, the Sn and Sp values were computed against the APPRIS annotation of genes of major protein isoforms.

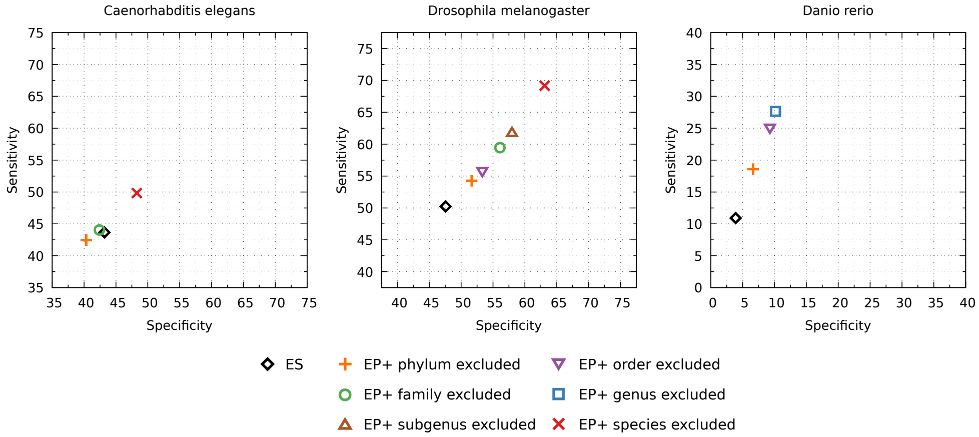


Figure S7: Same comparison as in Figure 5 in the main text, the accuracy is computed against the APPRIS annotation of genes of major protein isoforms.

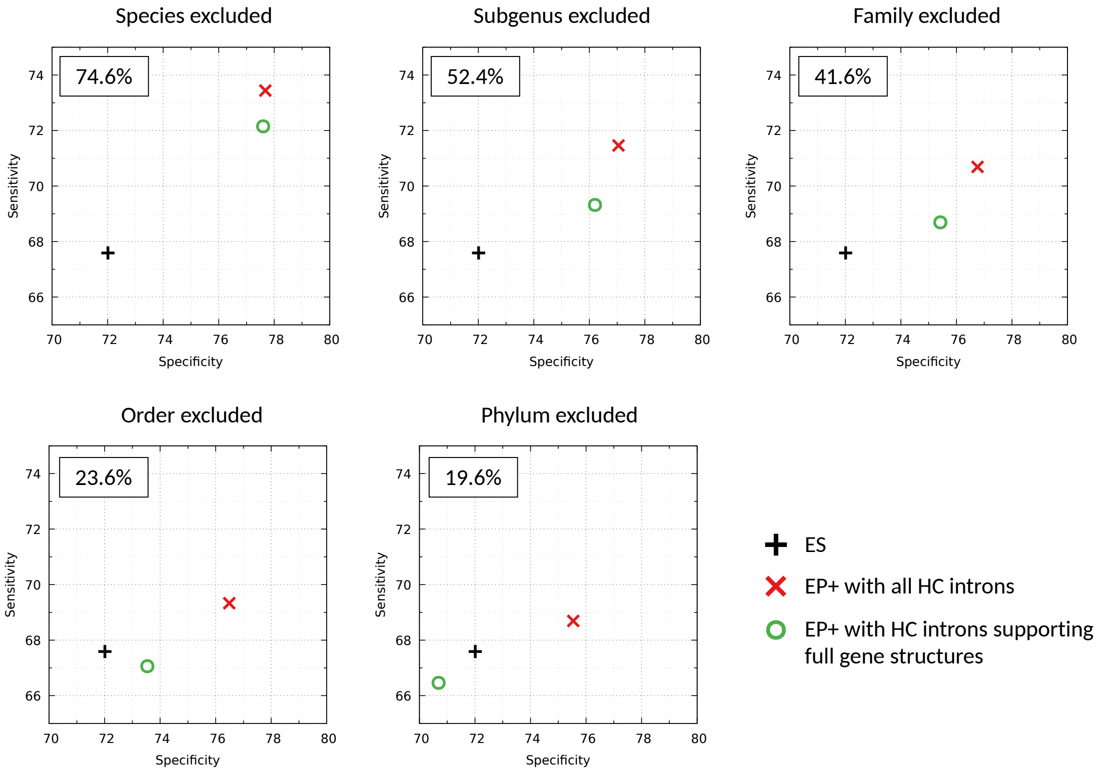


Figure S8: Comparison of exon level accuracy between three gene prediction modes in *D. melanogaster*. Use of introns from incomplete gene alignments leads to significant increase in accuracy compared to using only introns from fully aligned gene structure. GeneMark-ES is represented by a plus symbol. GeneMark-EP+ used only High-Confidence (HC) introns and is represented by a red cross. GeneMark-EP+ represented by a green circle used a subset of HC introns. This subset corresponds to annotated gene structures with all the introns supported by HC introns. In each panel we show percentage of such introns among all HC introns.

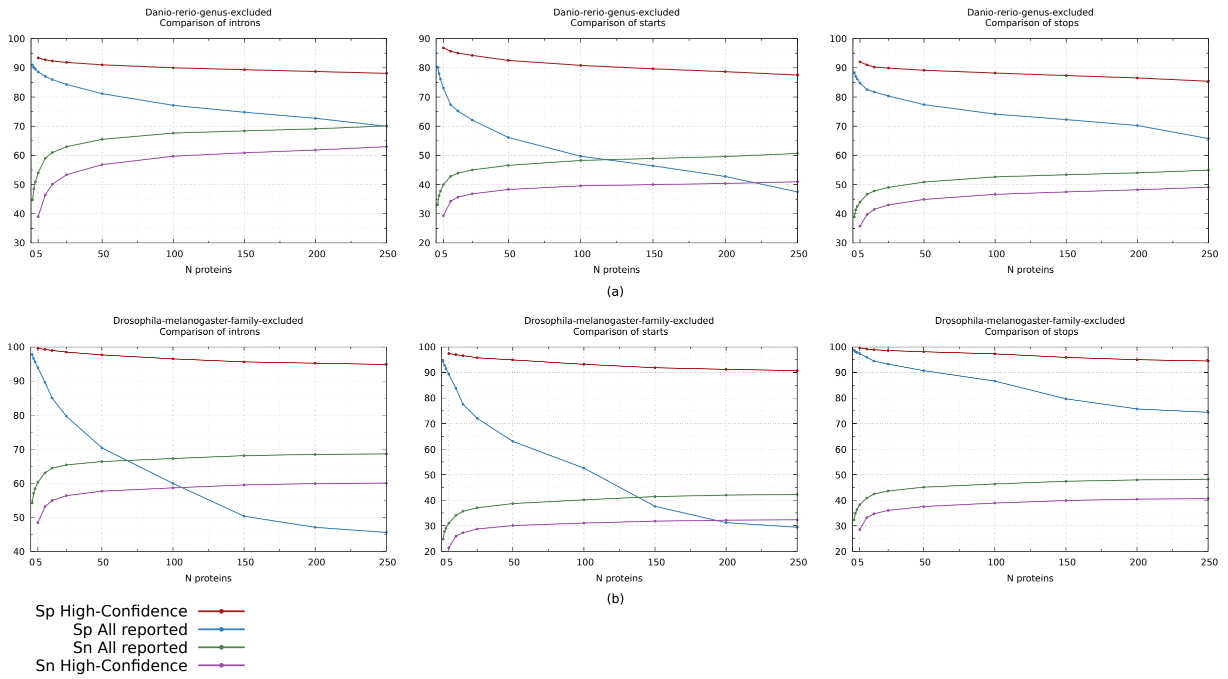


Figure S9: Effect of the maximum number of target proteins N per seed gene on sensitivity and specificity of hints to introns, start and stop codons. All reported and high-confidence hints are shown. Number N limits how many proteins found by DIAMOND and selected by the highest alignment scores are splice-aligned to a seed gene. Examples shown are (a) for large genome of *D. rerio*, and (b) for compact genome of *D. melanogaster*. The increase in Sn of intronic hints is larger in *D. rerio* because of a higher number of introns per gene (Table 1). Default value of N is set to 25 as a trade-off between computational speed of ProtHint and Sn of produced hints. Specificity of High-Confidence hints decreases slightly with increasing N. We recommend to use more strict (higher) SMC/IMC filtering thresholds when N $>$ 25 is selected.

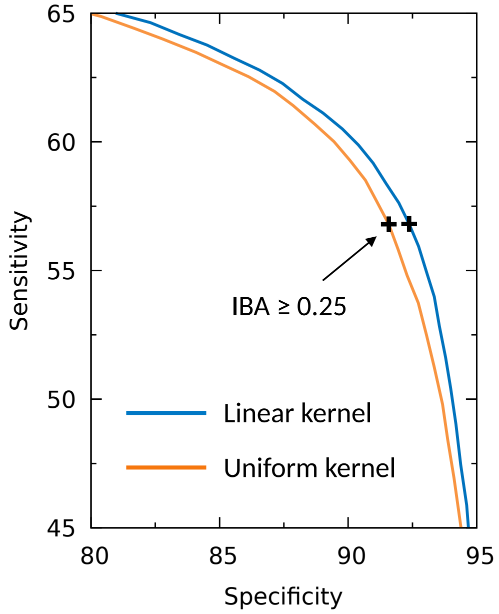


Figure S10: Advantage of the linear kernel. ProtHint intron hint Sp-Sn curves built for intron border alignment scores (IBA) computed with use of linear and uniform kernels (window width = 10). The crosses at the curves represent IBA score $\geq$ 0.25, with 0.25 being a value of the IBA threshold used for high-confidence intron selection. *D. melanogaster* genome with target proteins from species outside *Drosophilidae* family were used in this experiment.

#### Supplementary Tables

| Species | % of genes with a single protein isoform | # of isoforms per gene with multiple protein isoforms |
|---|---|---|
| *Caenorhabditis elegans* | 78% | 2.9 |
| *Drosophila melanogaster* | 74% | 3.3 |
| *Danio rerio* | 61% | 2.6 |

Table S1: Protein isoform statistics for species in APPRIS database.

| | | All reported starts | Filtered with SMC ≥ 4 | Filtered with SMC ≥ 4 and exon overlap = 0 |
|---|---|---|---|---|
| *N. crassa* | Sn | 65.3 | 39.4 | 38.8 |
| | Sp | 75.6 | 88.5 | **93.7** |
| *C. elegans* | Sn | 13.4 | 6.1 | 6.0 |
| | Sp | 68.2 | 94.5 | **95.8** |
| *A. thaliana* | Sn | 69.3 | 62.9 | 61.4 |
| | Sp | 70.9 | 89.8 | **94.4** |
| *D. melanogaster* | Sn | 37.7 | 29.6 | 29.2 |
| | Sp | 71.6 | 92.2 | **95.6** |
| *S. lycopersicum* | Sn | 48.9 | 43.6 | 42.8 |
| | Sp | 39.2 | 65.4 | **72.4** |
| *D. rerio* | Sn | 47.6 | 40.8 | 39.6 |
| | Sp | 61.4 | 80.8 | **84.1** |

Table S2: Sensitivity and specificity of ProtHint hints to gene starts for all the hints as well as for high-confidence hints. High specificity was achieved with filtering by start mapping coverage (SMC) scores as well as by removal of candidate starts overlapped by at least one target protein that suggested an alternative start upstream. Sensitivity was defined with respect to a full complement of starts, including starts of alternative isoforms. The tests were done in genus-excluded mode.

| | | | The level of exclusion of database proteins | | | | | | | | | | | |
|---|---|---|---|---|---|---|---|---|---|---|---|---|---|---|
| | | | Species | | Subgenus | | Genus | | Family | | Order | | Phylum | |
| ***N. crassa*** | ES | ET | | | | | EP | **EP+** | | | EP | **EP+** | EP | **EP+** |
| Gene Sn | 64.5 | 64.6 | | * | | * | 64.7 | **67.1** | | * | 64.3 | **66.1** | 64.1 | **64.7** |
| Gene Sp | 71.9 | 71.9 | | | | | 72.0 | **74.5** | | | 71.8 | **73.4** | 71.6 | **72.1** |
| Exon Sn | 75.7 | 75.9 | | | | | 75.7 | **77.8** | | | 75.4 | **77.2** | 75.3 | **75.8** |
| Exon Sp | 84.8 | 84.8 | | | | | 85.1 | **85.5** | | | 84.9 | **85.0** | 84.7 | **84.7** |
| Intron Sn | 79.5 | 79.8 | | | | | 79.6 | **82.5** | | | 79.3 | **82.0** | 79.1 | **80.3** |
| Intron Sp | 89.8 | 90.0 | | | | | 90.5 | **90.7** | | | 90.3 | **90.5** | 90.0 | **90.4** |
| ***C. elegans*** | ES | ET | EP | **EP+** | | | | | EP | **EP+** | | | EP | **EP+** |
| Gene Sn | 46.8 | 47.8 | 48.7 | **53.4** | | * | | * | 45.2 | **47.4** | | * | 43.5 | **45.7** |
| Gene Sp | 46.4 | 47.4 | 47.1 | **51.8** | | | | | 42.8 | **45.8** | | | 40.4 | **43.6** |
| Exon Sn | 81.0 | 81.0 | 81.3 | **82.4** | | | | | 80.0 | **80.3** | | | 79.6 | **79.9** |
| Exon Sp | 82.4 | 83.0 | 82.6 | **84.1** | | | | | 80.0 | **81.5** | | | 78.3 | **80.1** |
| Intron Sn | 87.5 | 87.3 | 87.5 | **88.4** | | | | | 86.4 | **86.6** | | | 86.1 | **86.3** |
| Intron Sp | 86.4 | 87.1 | 86.7 | **88.1** | | | | | 84.4 | **85.7** | | | 82.8 | **84.5** |
| ***A. thaliana*** | ES | ET | EP | **EP+** | | | EP | **EP+** | EP | **EP+** | EP | **EP+** | EP | **EP+** |
| Gene Sn | 55.8 | 57.2 | 57.5 | **73.7** | | * | 57.5 | **73.2** | 57.3 | **67.5** | 57.4 | **66.8** | 57.0 | **59.2** |
| Gene Sp | 54.0 | 55.3 | 55.4 | **69.4** | | | 55.4 | **69.1** | 55.3 | **64.6** | 55.4 | **64.0** | 55.3 | **57.3** |
| Exon Sn | 77.2 | 77.5 | 77.6 | **81.8** | | | 77.5 | **81.6** | 77.4 | **80.3** | 77.5 | **80.1** | 77.1 | **77.8** |
| Exon Sp | 79.2 | 80.4 | 80.5 | **84.8** | | | 80.5 | **84.7** | 80.6 | **83.7** | 80.6 | **83.5** | 80.6 | **81.4** |
| Intron Sn | 85.2 | 85.5 | 85.5 | **89.0** | | | 85.5 | **89.0** | 85.4 | **88.2** | 85.4 | **88.1** | 85.1 | **86.0** |
| Intron Sp | 82.4 | 83.9 | 83.9 | **87.7** | | | 83.9 | **87.7** | 83.9 | **87.1** | 84.0 | **87.0** | 84.1 | **85.1** |
| ***D. melanogaster*** | ES | ET | EP | **EP+** | EP | **EP+** | | | EP | **EP+** | EP | **EP+** | EP | **EP+** |
| Gene Sn | 50.2 | 52.4 | 53.3 | **69.2** | 52.9 | **61.8** | | * | 52.7 | **59.5** | 52.6 | **55.8** | 52.6 | **54.3** |
| Gene Sp | 47.6 | 48.8 | 50.0 | **63.1** | 49.6 | **58.0** | | | 49.6 | **56.1** | 50.1 | **53.3** | 49.7 | **51.7** |
| Exon Sn | 67.6 | 68.5 | 68.7 | **76.2** | 68.4 | **73.0** | | | 68.3 | **71.9** | 68.1 | **70.0** | 68.1 | **69.1** |
| Exon Sp | 72.0 | 73.6 | 74.8 | **80.9** | 74.5 | **78.9** | | | 74.6 | **78.2** | 75.1 | **77.0** | 74.8 | **76.0** |
| Intron Sn | 70.1 | 70.6 | 70.7 | **77.6** | 70.5 | **75.3** | | | 70.4 | **74.3** | 70.3 | **72.6** | 70.2 | **71.6** |
| Intron Sp | 75.5 | 77.3 | 78.7 | **84.2** | 78.5 | **82.9** | | | 78.6 | **82.3** | 79.2 | **81.5** | 78.8 | **80.3** |
| ***S. lycopersicum*** | ES | ET | | | | | EP | **EP+** | | | EP | **EP+** | EP | **EP+** |
| Gene Sn | 19.3 | 23.9 | | * | | * | 24.1 | **36.3** | | * | 24.2 | **33.5** | 24.2 | **26.1** |
| Gene Sp | 16.1 | 19.5 | | | | | 19.5 | **28.9** | | | 19.7 | **27.1** | 20.3 | **22.0** |
| Exon Sn | 65.2 | 68.8 | | | | | 69.0 | **75.5** | | | 68.9 | **74.3** | 68.0 | **69.5** |
| Exon Sp | 46.2 | 54.0 | | | | | 53.7 | **59.1** | | | 54.0 | **58.8** | 55.7 | **57.0** |
| Intron Sn | 71.9 | 76.3 | | | | | 76.3 | **84.6** | | | 76.2 | **83.5** | 75.5 | **77.8** |
| Intron Sp | 48.8 | 59.3 | | | | | 58.7 | **65.9** | | | 59.1 | **65.6** | 61.4 | **63.3** |
| ***D. rerio*** | ES | ET | | | | | EP | **EP+** | | | EP | **EP+** | EP | **EP+** |
| Gene Sn | 12.1 | 16.2 | | * | | * | 16.2 | **29.8** | | * | 16.2 | **27.0** | 16.4 | **20.4** |
| Gene Sp | 4.5 | 6.0 | | | | | 5.8 | **11.5** | | | 5.8 | **10.6** | 5.7 | **7.6** |
| Exon Sn | 64.0 | 66.5 | | | | | 66.5 | **72.7** | | | 66.3 | **71.6** | 66.1 | **68.3** |
| Exon Sp | 43.7 | 49.0 | | | | | 48.0 | **54.2** | | | 48.1 | **53.8** | 47.3 | **50.7** |
| Intron Sn | 63.7 | 66.2 | | | | | 66.3 | **73.1** | | | 66.2 | **72.1** | 66.0 | **68.7** |
| Intron Sp | 45.8 | 52.2 | | | | | 51.3 | **58.1** | | | 51.5 | **57.8** | 50.9 | **54.5** |

* See the first column to the right

Table S3: Comparison of GeneMark-ES, GeneMark-ET, GeneMark-EP and GeneMark-EP+ in terms of accuracy on gene, exon, and intron level. A gene was considered to be found if one of its annotated isoforms was predicted exactly. Exon and intron level Sn and Sp were defined with respect to a full complement of exons/introns, including ones from alternative isoforms. Gene level sensitivity for *D. rerio* was computed only with respect to complete genes. Accuracy of GeneMark-EP and GeneMark-EP+ determined with respect to annotation is shown for various types of protein database partition (species-excluded, etc).

| GeneMark- | ES | | EP+ genus excl. | | EP+ order excl. | | EP+ phylum excl. | |
|---|---|---|---|---|---|---|---|---|
| Test Set | **A** | B | **A** | B | **A** | B | **A** | B |
| Gene Sn | **22.7** | 4.8 | **53.0** | 8.4 | **47.8** | 8.0 | **34.0** | 6.9 |
| Exon Sn | **76.5** | 46.1 | **88.8** | 52.5 | **87.3** | 51.9 | **81.5** | 48.7 |
| Intron Sn | **79.5** | 53.3 | **93.9** | 61.7 | **92.7** | 60.9 | **86.3** | 56.8 |

| ProtHint HC | genus excluded | | order excluded | | phylum excluded | |
|---|---|---|---|---|---|---|
| Test Set | **A** | B | **A** | B | **A** | B |
| Intron Sn | **87.6** | 50.1 | **79.8** | 43.8 | **30.3** | 14.3 |
| Start Sn | **60.4** | 21.5 | **48.2** | 15.2 | **5.6** | 1.4 |
| Stop Sn | **69.3** | 20.5 | **54.3** | 14.7 | **8.1** | 1.7 |

Table S4: Accuracy assessment for *S. lycopersicum.* Only genes, which have all the introns in the gene supported by RNA-Seq mapping were selected into test set A and all the other genes were selected into set B. Single-exon genes were excluded from this analysis. RNA-Seq reads were mapped to genome by VARUS. Set A contained 15,832 genes with 84,424 introns. Set B contained 9,506 genes with 34,282 introns.

| | Original Annotation | Partial CDS removed, all transcripts | Complete transcripts | Incomplete transcripts | Complete genes | Incomplete genes |
|---|---|---|---|---|---|---|
| Exon Sn | 69.90 | 72.67 | **75.06** | 67.60 | 75.08 | 68.71 |
| Gene Sn | 23.98 | 24.34 | 27.11 | 0.19 | **29.84** | 12.11 |

Table S5: Comparison of GeneMark-EP+ predictions against full *D. rerio* annotation as well as annotation with *partial CDS* removed. Other columns show accuracy defined for a set of genes with complete/incomplete transcripts and for sets of complete/incomplete genes. A gene is considered complete if its transcripts are complete. All the numbers were generated in tests for protein database with proteins from species that belong to *D. rerio* genus excluded.

| | | The level of exclusion of database proteins | | | | | |
|---|---|---|---|---|---|---|---|
| Species | Genes in annotation | species | subgenus | genus | family | order | phylum |
| *N. crassa* | 10,785 | * | * | 9078 (84.2%) | * | 7,974 (73.9%) | 6,885 (63.8%) |
| *A. thaliana* | 27,445 | 23,854 (86.9%) | * | 23663 (86.2%) | 21,805 (79.4%) | 21,243 (77.4%) | 13,079 (47.7%) |
| *C. elegans* | 20,172 | 16,258 (80.6%) | * | * | 10,391 (51.5%) | * | 8,229 (40.8%) |
| *D. melanogaster* | 13,929 | 12,048 (86.5%) | 11,067 (79.5%) | * | 10,186 (73.1%) | 8,657 (62.2%) | 7,047 (50.6%) |
| *S. lycopersicum* | 33,562 | * | * | 23141 (69.0%) | * | 21,575 (64.3%) | 12,908 (38.5%) |
| *D. rerio* | 25,254 | * | * | 20439 (80.9%) | * | 19,809 (78.4%) | 14,856 (58.8%) |

* See the first column to the right

(a) All hints

| | | The level of exclusion of database proteins | | | | | |
|---|---|---|---|---|---|---|---|
| Species | Genes in annotation | species | subgenus | genus | family | order | phylum |
| *N. crassa* | 10,785 | * | * | 7,691 (71.3%) | * | 7,331 (68.0%) | 4,469 (41.4%) |
| *A. thaliana* | 27,445 | 23,029 (83.9%) | * | 22,879 (83.4%) | 20,224 (73.7%) | 19,961 (72.7%) | 9,127 (33.3%) |
| *C. elegans* | 20,172 | 11,439 (56.7%) | * | * | 7,242 (35.9%) | * | 6,115 (30.3%) |
| *D. melanogaster* | 13,929 | 11,752 (84.4%) | 10,180 (73.1%) | * | 9,332 (67.0%) | 7,372 (52.9%) | 5,497 (39.5%) |
| *S. lycopersicum* | 33,562 | * | * | 21,971 (65.5%) | * | 20,209 (60.2%) | 8,716 (26.0%) |
| *D. rerio* | 25,254 | * | * | 19,637 (77.8%) | * | 18,716 (74.1%) | 12,411 (49.1%) |

* See the first column to the right

(b) High-Confidence hints

Table S6: Annotated genes containing at least (a) one ProtHint hint (b) one High-Confidence ProtHint hint. Numbers (%) in the six genomes.

| Species | Genes in annotation | VARUS Hints |
|---|---|---|
| *N. crassa* | 10,785 | 7,462 (69.2%) |
| *A. thaliana* | 27,445 | 19,043 (69.4%) |
| *C. elegans* | 20,172 | 18,134 (89.9%) |
| *D. melanogaster* | 13,929 | 10,714 (76.9%) |
| *S. lycopersicum* | 33,562 | 19,158 (57.1%) |
| *D. rerio* | 25,254 | 21,841 (86.5%) |

Table S7: Annotated genes with at least one VARUS hint. Numbers (%) in the six genomes

| | | The level of exclusion of database proteins | | | | | |
|---|---|---|---|---|---|---|---|
| Species | Genes in annotation | species | subgenus | genus | family | order | phylum |
| *N. crassa* | 10,785 | * | * | 1,330 (12.3%) | * | 2,029 (18.8%) | 2,490 (23.1%) |
| *A. thaliana* | 27,445 | 2,534 (9.2%) | * | 2,683 (9.8%) | 4,028 (14.7%) | 4,372 (15.9%) | 7,672 (28.0%) |
| *C. elegans* | 20,172 | 1,013 (5.0%) | * | * | 1,812 (9.0%) | * | 1,911 (9.5%) |
| *D. melanogaster* | 13,929 | 856 (6.1%) | 1,452 (10.4%) | * | 1,821 (13.1%) | 2,247 (16.1%) | 2,549 (18.3%) |
| *S. lycopersicum* | 33,562 | * | * | 8,205 (24.4%) | * | 9,347 (27.8%) | 13,292 (39.6%) |
| *D. rerio* | 25,254 | * | * | 1,626 (6.4%) | * | 1,823 (7.2%) | 2,872 (11.4%) |

* See the first column to the right

Table S8: Annotated genes having no ProtHint or VARUS hints at all. Numbers (%) in the six genomes.

| | | ES | EP | EP+ Introns | EP+ Starts / Stops | EP+ Full |
|---|---|---|---|---|---|---|
| *Neurospora crassa* | Gene Sn / Sp | 64.5 / 71.9 | 64.7 / 72.0 | 66.0 / 73.6 | 66.3 / 73.4 | 67.1 / 74.5 |
| | **Exon Sn / Sp** | **75.7 / 84.8** | **75.7 / 85.1** | **77.3 / 84.9** | **76.7 / 85.8** | **77.8 / 85.5** |
| | Initial Sn / Sp | 70.9 / 81.3 | 70.2 / 81.2 | 72.4 / 82.0 | 72.2 / 82.3 | 73.2 / 82.7 |
| | Internal Sn / Sp | 77.4 / 88.2 | 77.2 / 89.1 | 80.4 / 87.5 | 77.8 / 89.8 | 80.4 / 88.7 |
| | Terminal Sn / Sp | 79.0 / 89.5 | 79.2 / 89.8 | 79.9 / 89.1 | 80.2 / 90.3 | 80.5 / 89.8 |
| | Single Sn / Sp | 74.7 / 70.7 | 74.5 / 69.9 | 73.3 / 71.5 | 75.7 / 70.5 | 74.3 / 71.7 |
| | Intron Sn / Sp | 79.5 / 89.8 | 79.6 / 90.5 | 82.5 / 90.4 | 80.2 / 90.9 | 82.5 / 90.7 |
| | Start Sn / Sp | 76.2 / 83.2 | 76.0 / 82.8 | 76.8 / 83.7 | 77.5 / 83.8 | 77.8 / 84.3 |
| | Stop Sn / Sp | 85.8 / 92.0 | 86.0 / 92.1 | 86.4 / 92.6 | 86.8 / 92.4 | 86.9 / 92.7 |
| *Caenorhabditis elegans* | Gene Sn / Sp | 46.8 / 46.4 | 45.2 / 42.8 | 46.4 / 45.0 | 46.3 / 43.8 | 47.4 / 45.8 |
| | **Exon Sn / Sp** | **81.0 / 82.4** | **80.0 / 80.0** | **80.2 / 81.2** | **80.2 / 80.4** | **80.3 / 81.5** |
| | Initial Sn / Sp | 53.5 / 63.4 | 53.1 / 60.1 | 53.3 / 61.7 | 53.8 / 60.8 | 54.0 / 62.4 |
| | Internal Sn / Sp | 90.7 / 87.7 | 89.6 / 86.4 | 89.9 / 87.1 | 89.6 / 86.7 | 89.8 / 87.4 |
| | Terminal Sn / Sp | 73.6 / 77.2 | 72.6 / 72.8 | 72.6 / 74.5 | 73.1 / 73.2 | 73.0 / 74.7 |
| | Single Sn / Sp | 15.6 / 50.5 | 16.6 / 46.5 | 16.7 / 48.3 | 18.1 / 47.4 | 17.8 / 48.8 |
| | Intron Sn / Sp | 87.5 / 86.4 | 86.4 / 84.4 | 86.7 / 85.5 | 86.4 / 84.7 | 86.6 / 85.7 |
| | Start Sn / Sp | 53.7 / 64.8 | 53.4 / 61.5 | 53.6 / 63.2 | 54.0 / 62.3 | 54.3 / 63.7 |
| | Stop Sn / Sp | 73.5 / 78.0 | 72.6 / 73.5 | 72.6 / 75.3 | 73.1 / 73.9 | 73.0 / 75.4 |
| *Arabidopsis thaliana* | Gene Sn / Sp | 55.8 / 54.0 | 57.5 / 55.4 | 65.2 / 63.1 | 65.7 / 61.4 | 73.2 / 69.1 |
| | **Exon Sn / Sp** | **77.2 / 79.2** | **77.5 / 80.5** | **80.1 / 82.5** | **79.6 / 83.0** | **81.6 / 84.7** |
| | Initial Sn / Sp | 60.5 / 68.9 | 61.1 / 69.5 | 63.3 / 71.4 | 66.7 / 73.9 | 67.9 / 75.3 |
| | Internal Sn / Sp | 87.1 / 83.4 | 87.3 / 85.1 | 90.6 / 87.1 | 87.6 / 87.3 | 90.5 / 89.2 |
| | Terminal Sn / Sp | 61.2 / 72.2 | 61.9 / 72.9 | 63.7 / 74.6 | 66.3 / 76.0 | 66.9 / 76.9 |
| | Single Sn / Sp | 58.6 / 74.2 | 59.1 / 73.3 | 58.3 / 76.1 | 64.3 / 75.9 | 63.7 / 78.5 |
| | Intron Sn / Sp | 85.2 / 82.4 | 85.5 / 83.9 | 89.0 / 86.3 | 86.0 / 85.5 | 89.0 / 87.7 |
| | Start Sn / Sp | 65.4 / 74.1 | 65.8 / 74.1 | 66.5 / 75.3 | 71.4 / 78.0 | 71.3 / 78.7 |
| | Stop Sn / Sp | 67.0 / 77.0 | 67.8 / 77.5 | 68.4 / 78.8 | 71.8 / 79.5 | 71.7 / 80.2 |
| *Drosophila melanogaster* | Gene Sn / Sp | 50.2 / 47.6 | 52.7 / 49.6 | 55.4 / 53.4 | 57.0 / 52.5 | 59.5 / 56.1 |
| | **Exon Sn / Sp** | **67.6 / 72.0** | **68.3 / 74.6** | **70.9 / 76.6** | **69.8 / 76.3** | **71.9 / 78.1** |
| | Initial Sn / Sp | 55.0 / 59.8 | 56.3 / 61.4 | 57.4 / 63.8 | 59.8 / 64.0 | 60.2 / 66.1 |
| | Internal Sn / Sp | 75.9 / 78.0 | 76.0 / 82.0 | 80.1 / 83.2 | 76.1 / 83.8 | 79.8 / 85.0 |
| | Terminal Sn / Sp | 63.1 / 68.2 | 64.4 / 69.6 | 65.5 / 72.4 | 67.2 / 71.5 | 67.7 / 73.7 |
| | Single Sn / Sp | 50.8 / 73.4 | 52.7 / 71.7 | 51.9 / 73.1 | 55.7 / 71.6 | 54.8 / 72.6 |
| | Intron Sn / Sp | 70.1 / 75.5 | 70.4 / 78.6 | 74.3 / 81.1 | 71.0 / 79.9 | 74.3 / 82.3 |
| | Start Sn / Sp | 58.4 / 65.5 | 59.7 / 66.6 | 60.1 / 68.5 | 63.3 / 68.9 | 63.1 / 70.4 |
| | Stop Sn / Sp | 68.5 / 75.3 | 69.5 / 75.9 | 70.0 / 78.2 | 72.1 / 76.9 | 72.0 / 78.7 |
| *Solanum lycopersicum* | Gene Sn / Sp | 19.3 / 16.1 | 24.1 / 19.5 | 31.3 / 25.7 | 28.9 / 22.4 | 36.3 / 28.9 |
| | **Exon Sn / Sp** | **65.2 / 46.2** | **69.0 / 53.7** | **73.9 / 57.8** | **71.6 / 55.4** | **75.5 / 59.1** |
| | Initial Sn / Sp | 40.2 / 31.1 | 44.1 / 33.7 | 47.0 / 36.4 | 50.7 / 36.9 | 51.8 / 38.8 |
| | Internal Sn / Sp | 79.0 / 51.4 | 82.3 / 62.9 | 88.5 / 67.5 | 82.7 / 64.6 | 88.4 / 69.1 |
| | Terminal Sn / Sp | 49.7 / 37.5 | 55.1 / 41.1 | 59.1 / 44.7 | 61.4 / 43.8 | 62.9 / 46.2 |
| | Single Sn / Sp | 29.7 / 46.9 | 33.2 / 42.1 | 32.7 / 43.6 | 37.1 / 43.9 | 36.6 / 45.2 |
| | Intron Sn / Sp | 71.9 / 48.8 | 76.3 / 58.7 | 84.6 / 65.1 | 77.2 / 59.7 | 84.6 / 65.9 |
| | Start Sn / Sp | 44.4 / 39.2 | 47.9 / 40.5 | 49.5 / 42.7 | 54.5 / 43.8 | 54.6 / 45.3 |
| | Stop Sn / Sp | 53.9 / 46.7 | 58.2 / 48.4 | 60.4 / 51.1 | 63.9 / 50.6 | 64.2 / 52.3 |
| *Danio rerio* | Gene Sn / Sp | 12.1 / 4.5 | 16.2 / 5.8 | 21.8 / 8.6 | 24.4 / 8.5 | 29.8 / 11.5 |
| | **Exon Sn / Sp** | **64.0 / 43.7** | **66.5 / 48.0** | **71.2 / 52.4** | **68.8 / 50.3** | **72.7 / 54.2** |
| | Initial Sn / Sp | 29.3 / 15.4 | 34.3 / 17.4 | 37.7 / 20.9 | 45.6 / 22.8 | 47.0 / 25.7 |
| | Internal Sn / Sp | 71.3 / 52.7 | 73.2 / 59.4 | 78.3 / 63.0 | 73.6 / 61.3 | 78.2 / 64.6 |
| | Terminal Sn / Sp | 43.8 / 23.2 | 47.9 / 24.5 | 51.4 / 28.8 | 55.5 / 28.1 | 56.5 / 31.2 |
| | Single Sn / Sp | 32.9 / 26.0 | 37.3 / 23.3 | 38.0 / 25.4 | 50.3 / 26.1 | 50.1 / 27.8 |
| | Intron Sn / Sp | 63.7 / 45.8 | 66.3 / 51.3 | 73.0 / 57.0 | 67.0 / 52.9 | 73.1 / 58.1 |
| | Start Sn / Sp | 32.9 / 17.5 | 38.3 / 19.5 | 40.9 / 22.8 | 51.3 / 25.5 | 51.7 / 28.1 |
| | Stop Sn / Sp | 48.7 / 26.0 | 52.6 / 26.9 | 55.6 / 31.0 | 61.7 / 30.8 | 62.0 / 33.8 |

Table S9: Assessment of accuracy of GeneMark-ES, GeneMark-EP and GeneMark-EP+. GeneMark-EP+ was run with enforcement of (a) only high confidence intron hints, (b) only high confidence hints to gene starts and stops (c) enforcement of both (a) and (b). Accuracy is shown at gene level, exon level (for all exons and separately for initial, internal, terminal, and single exons), intron level as well as for starts and stops. All the numbers were obtained for tests in genus-excluded mode. A gene was considered to be found if one of its annotated isoforms was predicted exactly. Exon, start, stop and intron level Sn and Sp are defined with respect to a full complement of isoforms, including alternative types. Gene level sensitivity of *D. rerio* was computed only with respect to complete genes.

| | The level of exclusion of database proteins | | | | | | | | | | | |
|---|---|---|---|---|---|---|---|---|---|---|---|---|
| | Species | | Subgenus | | Genus | | Family | | Order | | Phylum | |
| ***N. crassa*** | | | | | All reported | High conf. | | | All reported | High conf. | All reported | High conf. |
| Intron Sn | | * | | * | 76.0 | 66.2 | | * | 69.9 | 60.7 | 60.2 | 37.3 |
| Intron Sp | | | | | 58.4 | **96.8** | | | 61.6 | **96.9** | 70.8 | **98.3** |
| Start Sn | | | | | 65.3 | 38.8 | | | 43.0 | 34.3 | 27.4 | 9.6 |
| Start Sp | | | | | 75.6 | **93.7** | | | 76.0 | **91.5** | 73.7 | **89.0** |
| Stop Sn | | | | | 65.7 | 40.0 | | | 44.1 | 35.9 | 29.6 | 10.9 |
| Stop Sp | | | | | 94.1 | **98.5** | | | 95.9 | **98.4** | 96.1 | **99.2** |
| ***C. elegans*** | All reported | High conf. | | | | | All reported | High conf. | | | All reported | High conf. |
| Intron Sn | 76.7 | 36.7 | | * | | * | 37.4 | 18.1 | | * | 26.0 | 12.9 |
| Intron Sp | 91.8 | **99.0** | | | | | 92.8 | **99.3** | | | 93.7 | **99.2** |
| Start Sn | 47.7 | 13.4 | | | | | 13.4 | 6.0 | | | 8.2 | 5.1 |
| Start Sp | 75.8 | **96.5** | | | | | 68.2 | **95.8** | | | 76.2 | **95.0** |
| Stop Sn | 54.8 | 18.1 | | | | | 18.9 | 8.8 | | | 10.8 | 7.3 |
| Stop Sp | 90.7 | **97.0** | | | | | 92.4 | **97.7** | | | 92.9 | **97.3** |
| ***A. thaliana*** | All reported | High conf. | | | All reported | High conf. | All reported | High conf. | All reported | High conf. | All reported | High conf. |
| Intron Sn | 88.4 | 85.0 | | * | 87.9 | 84.3 | 82.6 | 74.2 | 80.3 | 71.6 | 51.3 | 28.3 |
| Intron Sp | 85.8 | **97.3** | | | 86.0 | **97.5** | 90.9 | **98.8** | 91.2 | **98.8** | 95.0 | **99.6** |
| Start Sn | 71.1 | 62.0 | | | 69.3 | 61.4 | 52.8 | 39.4 | 46.7 | 37.8 | 9.9 | 4.0 |
| Start Sp | 69.9 | **94.4** | | | 70.9 | **94.4** | 78.2 | **94.8** | 77.6 | **94.4** | 54.2 | **93.1** |
| Stop Sn | 67.1 | 60.4 | | | 64.9 | 59.0 | 47.9 | 37.5 | 43.3 | 36.3 | 11.1 | 5.1 |
| Stop Sp | 88.6 | **95.1** | | | 89.6 | **95.4** | 94.4 | **97.4** | 94.4 | **97.4** | 94.1 | **99.1** |
| ***D. melanogaster*** | All reported | High conf. | All reported | High conf. | | | All reported | High conf. | All reported | High conf. | All reported | High conf. |
| Intron Sn | 79.8 | 74.6 | 72.8 | 62.6 | | * | 66.2 | 54.3 | 49.7 | 34.4 | 35.8 | 20.9 |
| Intron Sp | 83.5 | **98.9** | 79.6 | **98.8** | | | 79.5 | **98.8** | 80.5 | **99.0** | 88.4 | **99.5** |
| Start Sn | 70.3 | 60.7 | 49.8 | 36.5 | | | 37.7 | 29.2 | 22.3 | 15.9 | 14.1 | 9.7 |
| Start Sp | 79.5 | **97.4** | 75.6 | **96.7** | | | 71.6 | **95.6** | 73.4 | **94.5** | 75.0 | **93.5** |
| Stop Sn | 75.3 | 68.4 | 56.7 | 45.2 | | | 44.7 | 36.9 | 26.7 | 19.8 | 15.8 | 11.2 |
| Stop Sp | 94.8 | **99.3** | 94.2 | **98.8** | | | 92.8 | **98.5** | 94.5 | **98.9** | 95.8 | **99.2** |
| ***S. lycopersicum*** | | | | | All reported | High conf. | | | All reported | High conf. | All reported | High conf. |
| Intron Sn | | * | | * | 80.6 | 76.8 | | * | 76.0 | 69.4 | 46.4 | 25.7 |
| Intron Sp | | | | | 70.5 | **92.0** | | | 81.7 | **93.5** | 89.7 | **95.6** |
| Start Sn | | | | | 48.9 | 42.8 | | | 39.9 | 32.9 | 8.5 | 3.4 |
| Start Sp | | | | | 39.2 | **72.4** | | | 43.8 | **74.6** | 40.7 | **77.9** |
| Stop Sn | | | | | 51.9 | 46.6 | | | 42.3 | 35.6 | 10.1 | 4.9 |
| Stop Sp | | | | | 69.9 | **83.6** | | | 76.9 | **85.5** | 85.8 | **92.0** |
| ***D. rerio*** | | | | | All reported | High conf. | | | All reported | High conf. | All reported | High conf. |
| Intron Sn | | * | | * | 65.5 | 55.8 | | * | 61.2 | 50.1 | 37.6 | 24.3 |
| Intron Sp | | | | | 84.4 | **92.2** | | | 86.8 | **93.5** | 90.1 | **96.8** |
| Start Sn | | | | | 47.6 | 39.6 | | | 39.6 | 31.3 | 14.3 | 8.7 |
| Start Sp | | | | | 61.4 | **84.1** | | | 70.4 | **85.7** | 64.1 | **89.5** |
| Stop Sn | | | | | 52.1 | 46.3 | | | 46.2 | 38.9 | 17.3 | 11.2 |
| Stop Sp | | | | | 79.8 | **89.8** | | | 85.6 | **91.8** | 87.6 | **95.5** |

* See the first column to the right

Table S10: Performance of ProtHint: Sensitivity and specificity of hints to introns, gene start and stop codons. Some cells of the table are left empty due to a low number or even complete absence of species within particular taxonomic ranks (Table 2). The results are shown for *all reported* hints as well as *high-confidence* hints. The accuracy is computed based on genome annotation including annotation of alternative isoforms.

| Species | Introns in the APPRIS set of principal isoforms | | |
|---|---|---|---|
| | All | In regions coding for conserved domains | |
| *D. melanogaster* | 41,010 | 21,562 | (52.6%) |
| *C. elegans* | 102,254 | 50,134 | (49.0%) |
| *D. rerio* | 178,867 | 106,288 | (59.4%) |

Table S11: Numbers of all annotated introns in the APPRIS set of principal isoforms and numbers of introns located within regions encoding conserved protein domains.

| Species | Exclusion level | High-confidence introns matching APPRIS introns | | | All reported introns matching APPRIS introns | | |
|---|---|---|---|---|---|---|---|
| | | All | In domains | | All | In domains | |
| *Drosophila melanogaster* | Species | 33,894 | 18,934 | (55.9%) | 35,338 | 19,414 | (54.9%) |
| | Subgenus | 28,437 | 17,475 | (61.5%) | 32,413 | 18,917 | (58.4%) |
| | Family | 24,670 | 16,057 | (65.1%) | 29,576 | 18,257 | (61.7%) |
| | Order | 15,829 | 11,984 | (75.7%) | 22,620 | 16,016 | (70.8%) |
| | Phylum | 9,719 | 8,222 | (84.6%) | 16,535 | 13,110 | (79.3%) |
| *Caenorhabditis elegans* | Species | 38,912 | 30,346 | (78.0%) | 80,402 | 45,210 | (56.2%) |
| | Family | 19,155 | 16,556 | (86.4%) | 39,379 | 29,270 | (74.3%) |
| | Phylum | 13,668 | 12,216 | (89.4%) | 27,464 | 23,140 | (84.3%) |
| *Danio rerio* | Genus | 108,236 | 71,239 | (65.8%) | 126,010 | 80,307 | (63.7%) |
| | Order | 97,457 | 67,335 | (69.1%) | 118,131 | 78,078 | (66.1%) |
| | Phylum | 47,860 | 40,117 | (83.8%) | 73,568 | 58,355 | (79.3%) |

Table S12: Change in fraction of high-confidence and all reported intron hints mapped to conserved protein domains when protein database size is changed from largest (species or genus excluded) to smallest (phylum excluded). Gene annotation is taken from APPRIS database.
